# TSTScope Unifies Single-Cell Multi-Omics to Identify Functional T Cell States Predictive of Immunotherapy Response

**DOI:** 10.64898/2026.01.08.698283

**Authors:** Shiwei Cao, Jinyu Cheng, Fengao Wang, Chenxin Yi, Jiajun Chen, Keyue Wang, Lulu Liu, Junwei Liu, Yixue Li

## Abstract

Immune checkpoint blockade (ICB) can produce durable responses in cancer, but reliable predictors of benefit are still lacking. CD8⁺ tumor-specific T cells are essential for ICB efficacy, yet it remains unclear which functional states of these cells determine therapeutic success. To address this, we developed TSTScope, an interpretable deep learning framework that integrates single-cell transcriptomic and T-cell receptor sequencing data to generate unified representations of CD8⁺ T-cell identity. By applying TSTScope to non-small cell lung cancer (NSCLC) datasets, we characterized the gene programs defining tumor specificity and computationally inferred a population of potential TSTs. Crucially, we demonstrate that clinical response is not a product of TST abundance, but is instead governed by their functional state. We derived the MPR score, a metric capturing this functional potential, which proved to be a robust predictor of treatment outcomes. In an independent validation cohort, the MPR score significantly outperformed established biomarkers. Collectively, TSTScope identifies a distinct functional state of tumor-specific T cells as a primary determinant of ICB efficacy, providing both a mechanistic framework and a potent tool for precision immunotherapy.

## INTRODUCTION

Immune checkpoint blockade (ICB) has reshaped cancer treatment, offering durable responses in a subset of patients. However, the majority fail to benefit, highlighting the critical need for reliable biomarkers to inform treatment decisions^1^. Accumulating evidence positions CD8⁺ tumor-specific T cells (TSTs) as central mediators of ICB efficacy, given their recognition of neoantigens through T cell receptors (TCRs)^2^. However, the specific cellular states that determine whether TSTs can mount an effective anti-tumor response remain poorly defined.

Recent advances in single-cell technologies, particularly scRNA-seq and scTCR-seq, now enable high-resolution profiling of T-cell states and clonality. scRNA-seq captures transcriptional heterogeneity, while scTCR-seq yields clonal and antigen-specific insights. Despite these advances, integrating these modalities to identify tumor-reactive T cells and correlate their functional states with clinical outcomes remains a major challenge. Existing approaches, including mvTCR^3^, scNAT^4^, UniTCR^5^, and MIST^6^, utilize concatenate or contrastive learning to align multi-modal data, but fail to capture the inherent interplay between antigen recognition and transcriptional programs. Furthermore, raw TCR embeddings are often confounded by germline V/J gene usage, masking CDR3-driven specificity signals^7^. Additionally, the limited interpretability of existing models further hampers their translation to clinical biomarker development^8,9^.

Consequently, consistent TST identification across heterogeneous datasets remains elusive. Prior strategies have depended on validated gene signatures, including established markers (CXCL13 and ENTPD1/CD39)^10^ or predefined TST gene sets, as in Oliviera et al.^11^, Lowery et al. (NeoTCR)^12^, and Yossef et al. (NeoTCRPBL)^13^. These signatures, however, are prone to batch effects and lack the generalizability of these gene expressions for cross-study predictions. Additionally, machine learning tools like MANAscore have sought to infer potential TSTs (pTSTs) across datasets using known signatures^14^. Nevertheless, these approaches remain constrained by their reliance on transcriptomic data alone, leaving the integration of transcriptional features with antigen specificity from TCR sequences largely unexplored. Furthermore, characterizing the functional heterogeneity of TSTs poses a secondary challenge^15,16^. Although stem-like precursor exhausted T cells (Tpex) are recognized as pivotal for sustaining ICB responses, a consensus on the precise definition of the Tpex population remains lacking^17,18^. Current analytical frameworks struggle to effectively resolve TST heterogeneity or elucidate the dynamic trajectories driving the transition from effector states to dysfunction. In particular, there is a distinct lack of quantitative metrics to measure the “effector potential” of TSTs, which is vital for predicting patient outcomes in the context of ICB therapy^19,20^.

To address these limitations, we introduce TSTScope, a multimodal deep learning framework that jointly integrates scRNA-seq and scTCR-seq data. TSTScope utilizes a gene program (GP)-constrained variational autoencoder (VAE) to construct an interpretable latent space, effectively unifying antigen specificity with transcriptional states. This approach enables both the robust identification of TSTs and high-resolution mapping of their functional heterogeneity. Applying TSTScope to non-small cell lung cancer (NSCLC) cohorts, we reveal that the functional context of TSTs, rather than mere cellular abundance, is the primary driver of ICB efficacy. Leveraging these insights, we developed the “MPR score,” a novel metric designed to quantify the effector potential of TSTs. The MPR score significantly outperforms existing biomarkers, achieving an AUC of 0.76 in independent validation cohorts. This establishes a robust, quantifiable link between TST heterogeneity and clinical outcomes, providing a superior predictive tool for immunotherapy response.

## RESULTS

### Model framework of TSTScope

To address the need for an integrated model combining scRNA-seq and scTCR-seq data for functional analysis of TSTs, TSTScope was developed to generate a unified, interpretable representation of T cells by merging transcriptomic profiles with paired TCR repertoires (Fig. 1A). Specifically, TSTScope employs a GP-constrained VAE framework to embed scRNA-seq data into a 50-dimensional latent space, where each dimension is explicitly constrained to represent a biologically meaningful T-cell GP, such as naïve, effector memory, or exhausted states. Concurrently, single-chain β TCR sequences from paired T cells are embedded using a fine-tuned TCR-BERT model^21^ and projected into the same latent space through fully connected layers (Methods). To capture the interplay between transcriptional state and TCR specificity, TCR and gene embeddings are aligned through a mixed alignment module prior to integration at clonotype level (Methods). This strategy preserves modality-specific information while capturing cross-modal interactions. Latent space interpretability is ensured by the VAE decoder, which reconstructs gene expression solely from the genes within each GP, enforcing each latent variable to correspond to specific biological concept^22^. Additionally, a TCR decoder reconstructs TCR embeddings from the latent space, maintaining fidelity to TCR information, and the joint embeddings from both gene and TCR were used for downstream analysis (Methods).

**Fig. 1.**
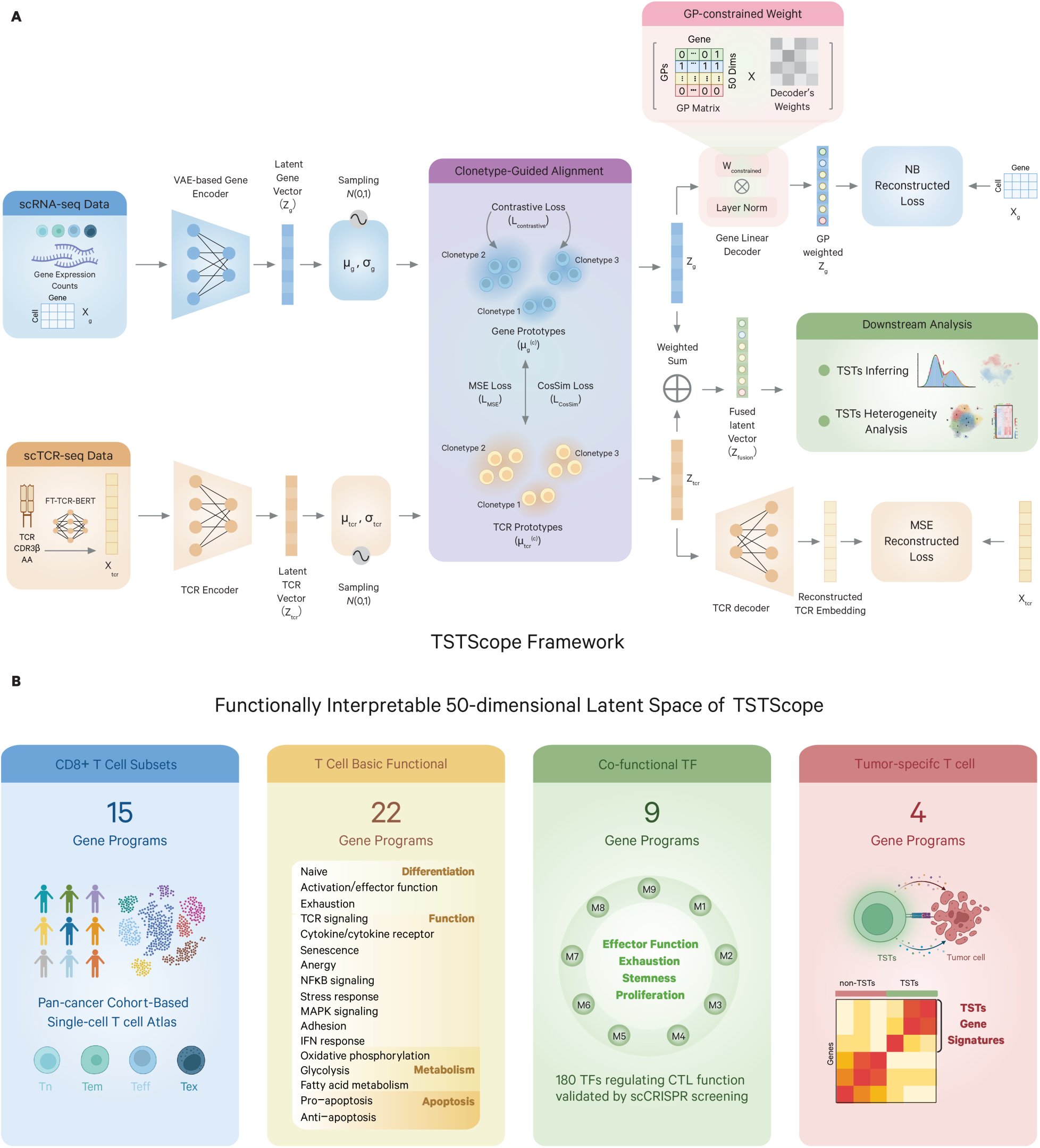
The TSTScope Framework. (**A**) Overview of the TSTScope architecture integrating scRNA-seq and paired scTCR-seq data into a unified latent representation. (**B**) Functionally interpretable 50-dimensional latent space of TSTScope organized by four categories of curated T-cell gene programs.

To enable comprehensive representation of T-cell functions, particularly for TST cells, the latent space is hierarchically organized around 50 curated T-cell GPs capturing cellular states from four complementary perspectives (Fig. 1B). These include: (i) T-cell subset programs (15 GPs) representing canonical CD8⁺ T subsets across their life cycle^23^; (ii) Core functional programs (22 GPs) covering essential processes such as TCR signaling, cytokine responses, metabolism, proliferation, and apoptosis^24,25^; (iii) Co-functional transcription factor modules (9 GPs) reflecting regulatory circuits of stemness, effector function, proliferation, and exhaustion, derived from large-scale CRISPR screens^26^; and (iv) Tumor-specific programs (4 GPs) distinguishing TSTs from bystanders, curated from validated datasets^13,27–29^ (Table S1). The combined gene list from all GPs was then used for further model building and training (Methods). This structure ensures biologically consistent cross-modal representations, minimizes batch effects, and provides a robust, biologically grounded latent space for analyzing T-cell diversity.

### Joint learning resolves TCR antigen specificity

We evaluated whether TSTScope’s joint learning strategy produces biologically meaningful TCR embeddings using a publicly available 10X Genomic dataset with known antigen labels (Methods). We compared the joint learned TSTScope’s TCR embeddings with transcriptomic context to its initial embeddings derived from the fine-tuned TCR-BERT model^21^. Additionally, we benchmarked these against TCR embeddings inferred from two pre-trained TCR sequence models, TCR-BERT^21^ and scCVC^7^. Uniform manifold approximation and projection (UMAP) was used to visualize TCR embeddings and related antigen specificities (Fig. 2A). To quantitatively assess the intrinsic clustering structure of antigen-specific T cells, we applied K-means clustering to the embeddings and calculated the Normalized Mutual Information (NMI) and Adjusted Rand Index (ARI). Among all methods, TSTScope achieved the highest NMI (0.129) and ARI (0.05), indicating superior separation of antigen-specific populations in the unsupervised space.

**Fig. 2.**
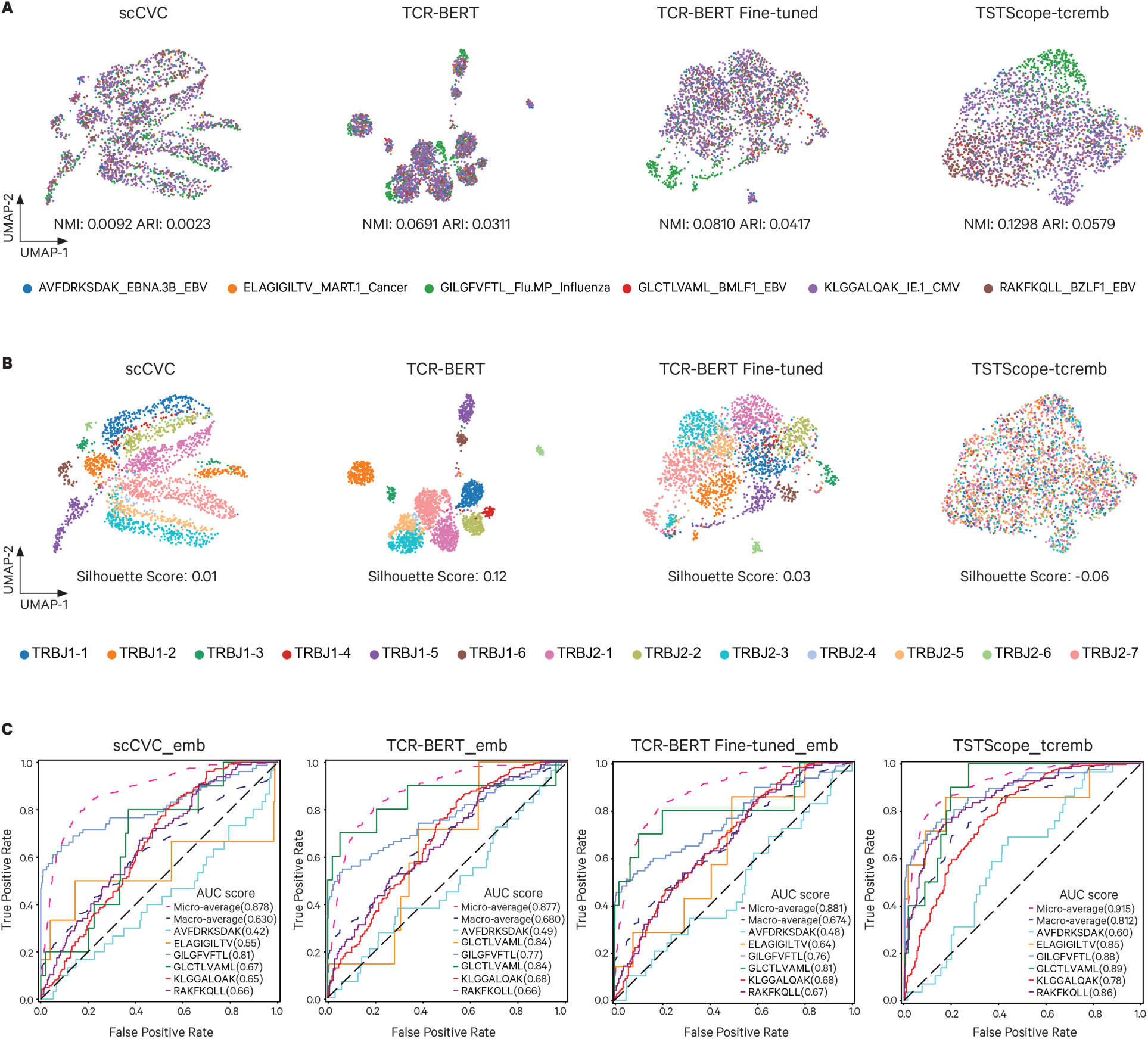
Joint learning improves biological relevance of TCR embeddings. (**A**) UMAP visualization of TCR embeddings inferred by scCVC, TCR-BERT, fine-tuned TCR-BERT, and TSTScope (n=2,406 cells). Points are colored by known antigen specificity. Performance is quantified by Normalized Mutual Information (NMI) and Adjusted Rand Index (ARI) scores, as indicated in each panel. (**B**) UMAP visualization of the same TCR embeddings as in (A), colored by TRBJ gene usage. The degree of clustering bias is quantified by the Silhouette score, as indicated in each panel. (**C**) Receiver operating characteristic (ROC) curves evaluating the ability of different TCR embeddings to predict antigen specificity. Each plot displays the Area Under the Curve (AUC) for individual antigens, alongside the micro- and macro-average AUC scores for a comprehensive performance comparison.

Furthermore, embeddings derived solely from TCR sequence were heavily confounded by conserved germline-encoded J gene segments, leading to clustering driven by germline similarity rather than antigen specificity (Fig. 2B). In contrast, TSTScope effectively distinguished TCRs based on their antigen targets, achieving the lowest Silhouette Score of −0.06 (Fig. 2B). To further evaluate the embeddings’ discriminative performance, we trained a multi-class classifier to predict T-cell antigen specificity (Methods). TSTScope’s joint TCR embedding achieved the highest accuracy, with a micro-average AUC of 0.915, surpassing the initial embedding (0.881), scCVC (0.878) and TCR-BERT (0.877) (Fig. 2C). Across multiple metrics, including antigen-specific accuracy, precision, recall, F1 score, and AUC scores, TSTScope consistently outperformed competing models (fig. S1A). These findings demonstrate that integrating transcriptional context effectively suppresses germline-associated artifacts and emphasizes the hypervariable CDR3 region.

### TSTScope accurately represents neoantigen-specific T cells

A primary objective of T-cell repertoire analysis in the Tumor Microenvironment (TME) is to identify T cells that recognize tumor-specific neoantigens. We next evaluated TSTScope’s ability to detect distinct transcriptomic and TCR patterns of TSTs. We utilized a NSCLC dataset from Caushi et al.^30^, comprising 16 patients who underwent paired scRNA-seq and scTCR-seq profiling. After quality control, 188,804 CD8^+^ T cells were retained for downstream analysis. Within this cohort, 2,083 TSTs and 1,420 non-TSTs cells from 9 patients were experimentally validated using the mutation-associated neoantigen functional expansion of specific T cells (MANAFEST) assay and ViralFEST, respectively. We visualized the distribution of these cells within the latent spaces generated by TSTScope and compared the results with those from alternative approaches.

We benchmarked TSTScope’s joint embeddings against three established transcriptome-only methods: principal component analysis (PCA) with Harmony batch correction^31^, a pre-trained cross-study human CD8^+^ T cell atlas and automatic subtype annotation scAtlasVAE^32^, and a parallel GP-constrained gene-expression model, ExpiMap, using same GPs as TSTScope^22,33^. This comparison was designed to assess the added value of integrating TCR sequence information with gene expression data, especially for the scAtlasVAE and ExpiMap models with prior biological information. UMAP visualizations and cell clustering analyses revealed that TSTScope achieved superior separation of TSTs, forming compact and distinct cell clusters. In contrast, the transcriptome-only methods yielded less-defined groupings with substantial intermingling of other CD8^+^ T cells (Fig. 3, A and B).

**Fig. 3.**
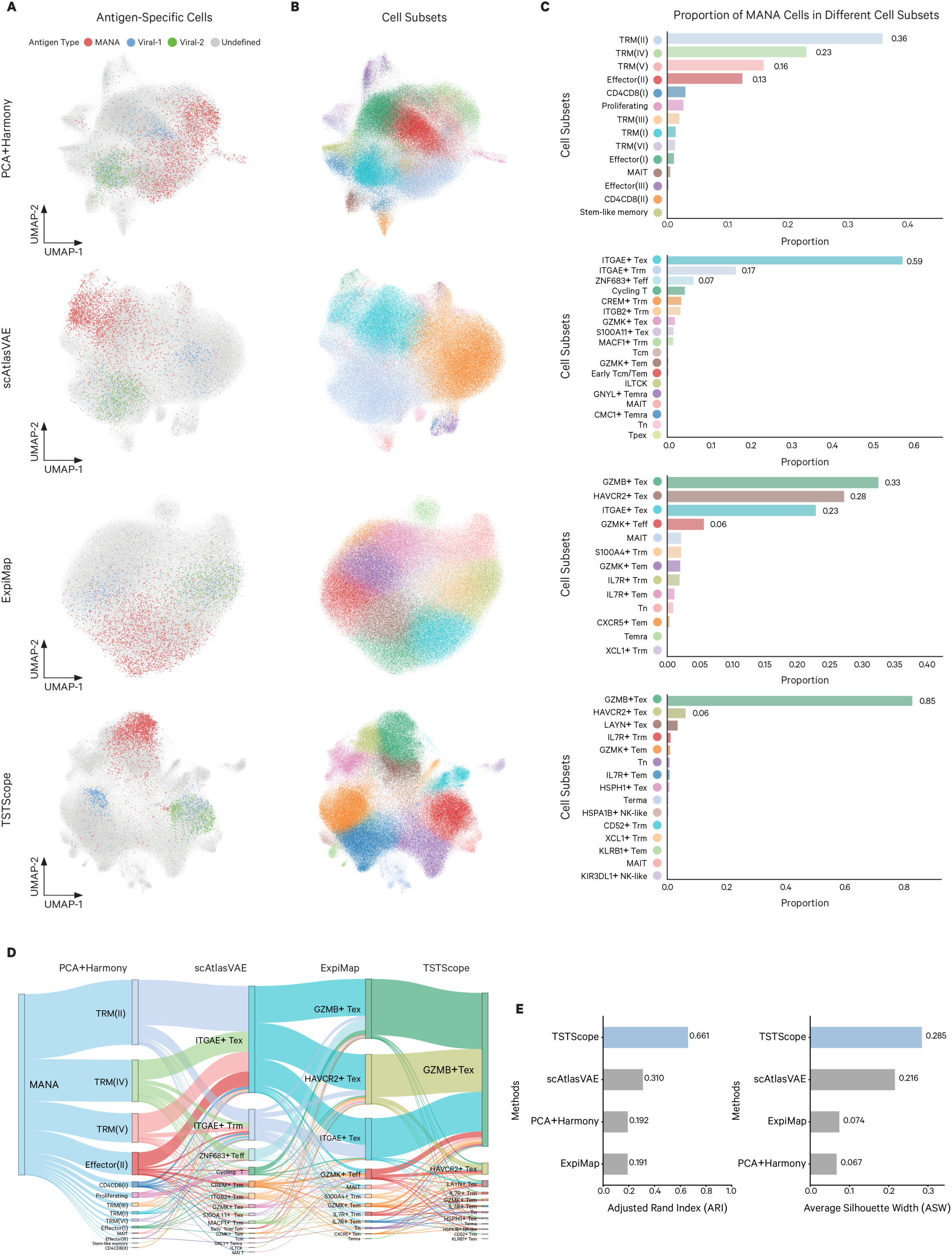
TSTScope enhances the identification and characterization of neoantigen-specific T cells. (**A-B**) UMAP visualizations of 188,804 CD8^+^ T cells from NSCLC patients (n=16), colored by antigen specificity validated by MANAFEST/ViralFEST assays (**A**) and cell subset annotations (**B**) across four integration methods: PCA+Harmony, scAtlasVAE, ExpiMap, and TSTScope. (**C**) Distribution of mutation-associated neoantigen (MANA)-specific cells across subsets identified by each method, numerical labels indicate proportions for subsets exceeding a 5% threshold. (**D**) Alluvial plot illustrating the mapping and concordance of validated MANA cells from original study clusters (left) through scAtlasVAE and ExpiMap to TSTScope-defined subsets (right). (**E**) Quantitative benchmarking of clustering performance across methods using Adjusted Rand Index (ARI; left) and Average Silhouette Width (ASW; right), calculated based on ground-truth antigen specificity labels.

We further compared the ability of different approaches to capture the TST enrichment at the cell-subset level. In the original result by Caushi et al., 14 CD8^+^ T-cell clusters were initially defined, while scAtlasVAE automatically assigned cells to 18 pre-defined cell subtypes. In contrast, ExpiMap and TSTScope model identified 10 and 15 T cell clusters, respectively, with annotations based on prior functional transcriptomic studies of T cells (Fig. 3B). Compositional analysis revealed that validated TSTs were predominantly enriched in tissue-resident memory (TRM)-like subsets with high exhaustion profiles, including TRM(II), TRM(IV), and TRM(V) from the original clustering (Fig. 3C). Similarly, scAtlasVAE assigned approximately 59% and 17% of TSTs to *ITGAE*^+^ terminally exhausted (Tex) and *ITGAE*^+^ Trm clusters, respectively. For both ExpiMap and TSTScope, the majority of TSTs were grouped into a *GZMB*^+^ Tex cluster characterized by cytotoxic potential and co-expression of exhaustion- and tissue-residency-related genes, alongside canonical tumor-specific T cell markers such as *CXCL13* and *ENTPD1* (fig. S2). Notably, TSTScope’s joint embedding mapped nearly 85% of TSTs to the *GZMB*^+^ Tex cluster, with an alluvial plot confirming strong concordance with experimentally validated TSTs (Fig. 3, C and D).

To quantitatively benchmark tumor-specific T-cell (TST) identification across different approaches, we performed a compositional analysis. The results demonstrated that clusters identified by TSTScope most consistently satisfied the dual criteria of high purity and broad population coverage. Consequently, TSTScope showed the strongest alignment with ground-truth labels, achieving superior performance metrics with an adjusted Rand index (ARI) of 0.661 and an average silhouette width (ASW) of 0.285 (Fig. 3E and fig. S1B). These findings suggest that integrating TCR sequence context with transcriptomic profiles significantly improves the resolution and accuracy of TST identification.

### Latent gene programs predict potential tumor-specific T cells

Leveraging the interpretable, GP-based latent space of TSTScope, we assessed the distribution of specific GPs to characterize potential TSTs (pTSTs). Enrichment analyses distinguished validated TSTs (MANA-specific T cells) from non-TSTs (Viral-specific T cells). Validated TSTs were preferentially enriched in GPs associated with T-cell exhaustion and tissue residency (e.g., OXPHOS_TEX, GZMK_TEX, TERM_TEX, ADHESION), as well as tumor-specific programs (e.g., TUMOR_SPECIFIC, TUMOR_REACTIVITY). Conversely, non-TSTs showed heightened activity in early memory programs (TCR_SIGNALING, TCM, EARLY_TEM) and stemness related programs (NAÏVE, MODULE_5) (Fig. 4, A and B). These patterns of gene-program activity were also consistently preserved within TST-associated subsets (GZMB+ Tex, HAVCR2+ Tex), and viral-specific T-cell-associated subsets (IL7R+ Trm, GZMK+ Tem) (fig. S3A and B), which align with established characterizations of TST biology^34–36^.

**Fig. 4.**
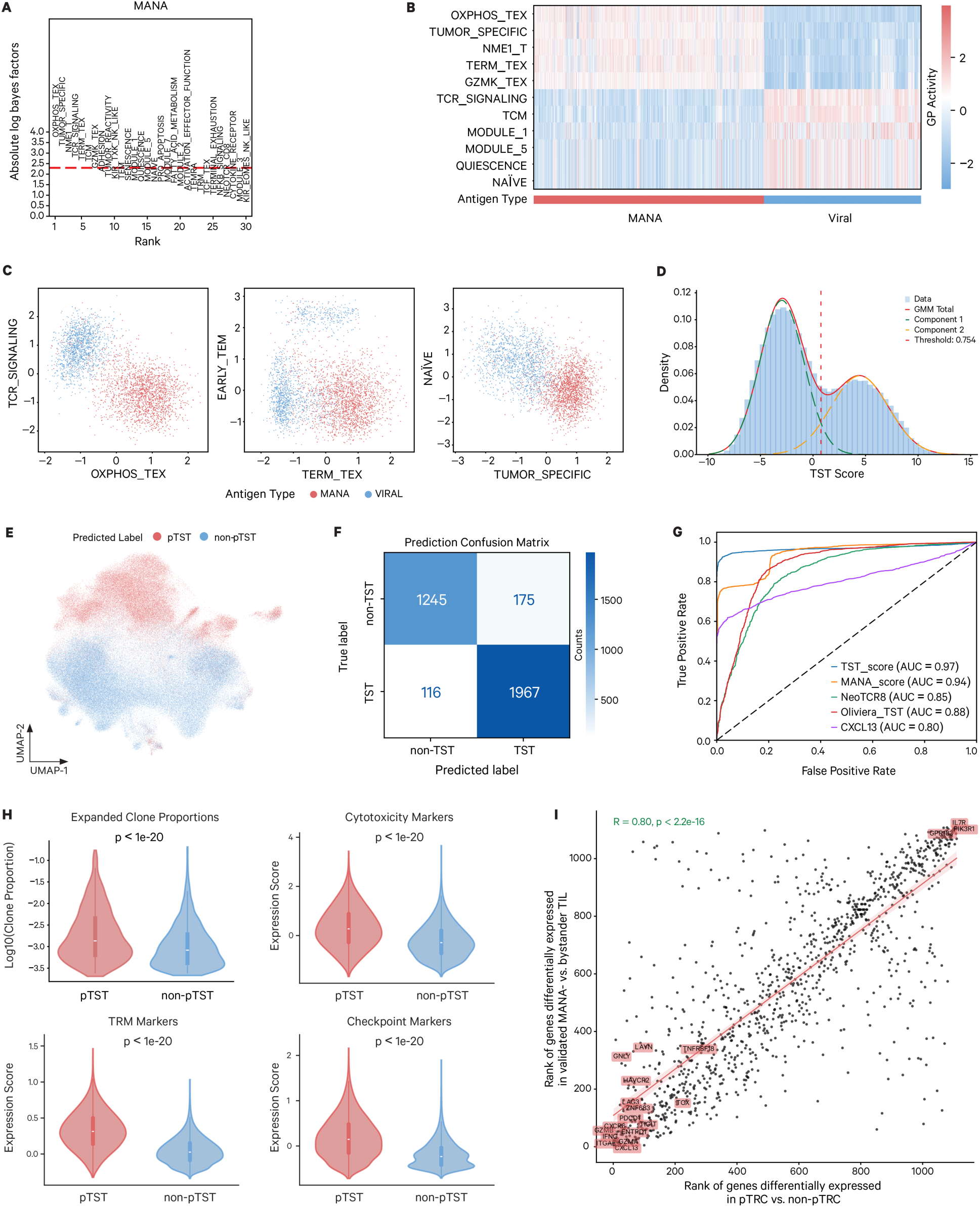
Latent gene programs and TST score accurately predict potential tumor-specific T cells. (**A-B**) Latent signature identification showing (**A**) the rank of gene programs (GPs) by absolute log Bayes factors and (**B**) heatmap of representative GP activities distinguishing validated MANA-specific (TST) from viral-specific (non-TST) cells. (**C**) Bivariate scatter plots of representative gene programs illustrating the robust transcriptional separation between tumor-specific and viral-specific cell populations. (**D**) Bimodal distribution of the TST score across the discovery cohort. A dual Gaussian Mixture Model (GMM) was used to adaptively define classification thresholds for potential TSTs (pTSTs). (**E**) UMAP projection of the Caushi’s NSCLC dataset (n=188,804 cells) colored by GMM-predicted pTST and non-pTST labels. (**F-G**) Performance evaluation of TSTScope via (**F**) confusion matrix and **(G)** ROC curve analysis benchmarking the TST score against established biomarkers and TST predictors. AUC scores of different biomarkers were labelled. (**H**) Comparison of predicted pTSTs and non-pTSTs across signatures for clonal expansion, cytotoxicity, tissue residency (TRM), and immune checkpoints. Statistical significance was determined by two-sided Mann-Whitney U tests and p-values were labelled. (**I**) Pearson correlation of differentially expressed gene (DEG) ranks between experimentally validated MANA-specific cells and TSTScope-predicted pTSTs. The correlation coefficient and p-value were labelled.

Across validated TSTs and TST-associated subsets, OXPHOS_TEX, TERM_TEX and TUMOR_SPECIFIC consistently ranked among the most robust positively enriched gene programs, whereas TCR_SIGNALING, EARLY_TEM and NAÏVE represented the most robustly depleted programs. The combined activity of positive and negative gene programs effectively separated validated TSTs from non-TSTs (Fig. 4C and fig. S3C). Additionally, among tumor-associated gene programs, TUMOR_SPECIFIC showed the strongest and most consistent enrichment across distinct TST groups, exceeding that of related programs such as TUMOR_REACTIVITY and NEOTCR_CD8 (fig. S3D). This underscores TSTScope’s ability to resolve subtle yet critical distinctions among overlapping gene signatures, thereby pinpointing a core transcriptional hallmark of TST identity.

Based on these insights, we defined a TST score as the combined activity of positively associated GPs (TUMOR_SPECIFIC, TERM_TEX, OXPHOS_TEX) minus that of negatively associated ones (TCR_SIGNALING, NAÏVE, EARLY_TEM). To identify pTSTs, we employed a dual Gaussian Mixture Model (GMM) to model the bimodal distribution of TST scores, distinguishing high-scoring pTSTs from the majority of low-scoring non-TSTs (Fig. 4D). The classification threshold was set at the intersection of the two Gaussian curves. To accommodate sample-specific variability, thresholds were derived adaptively for each sample, with a global cutoff applied only in the absence of bimodality (Methods). This data-driven approach eliminates arbitrary thresholds, ensuring robust and reproducible classification across datasets.

In the NSCLC dataset, the GMM-based classifier identified 74,128 pTSTs with high TST score among 188,804 CD8⁺ T cells. These pTSTs were enriched in subsets containing experimentally validated TSTs (Fig. 4E and fig. S3E), among which, 76.7% originated from in tumor tissue and were enriched within subsets that also contained experimentally validated TSTs (fig. S3F). Confusion matrix analysis revealed strong concordance with ground-truth labels, achieving a precision of 93.92% and specificity of 94.30% (Fig. 4F and fig. S3G). Notably, the TST score outperformed established biomarkers, including *CXCL13* expression, TST gene sets (NeoTCR8, Oliveira_TST), and the MANAscore predictor^14,27,28^, for pTSTs identification (Fig. 4G). The inferred pTSTs exhibited hallmark TST features, including enhanced clonal expansion and elevated expression of genes associated with tissue residency (*ITGAE*, *ZNF683*), cytotoxicity (*GZMB*, *GZMA*), and immune checkpoints (*LAG3*, *ENTPD1*, *TIGIT*) compared to non-pTST cells (Fig. 4H). The differential expression gene (DEG) analysis also confirmed strong correlation with validated TST profiles (Fig. 4I).

### Functional potential of pTSTs underlies ICB efficacy

To investigate the relationship between pTSTs and clinical outcomes, we analyzed their abundance and functional states in tumor tissue across ICB response groups in Caushi et al. dataset. This cohort include tumor sample from 15 patients: 6 with major pathological response (MPR) and 9 with non-MPR. Notably, neither the proportion of pTSTs in tumor tissue nor their average TST score showed a positive correlation with MPR (Fig. 5A). In fact, patients with MPR exhibited a trend toward lower average TST score, suggesting that the mere presence of tumor-reactive T cells is insufficient for clinical benefit. Instead, a high TST score may reflect terminally differentiated states that are refractory to reactivation. This interpretation was further supported by differential gene expression ranking analysis, which showed that pTSTs derived from MPR patients exhibited an overall transcriptional program largely opposite to that observed in validated TSTs (Fig. 5B).

**Fig. 5.**
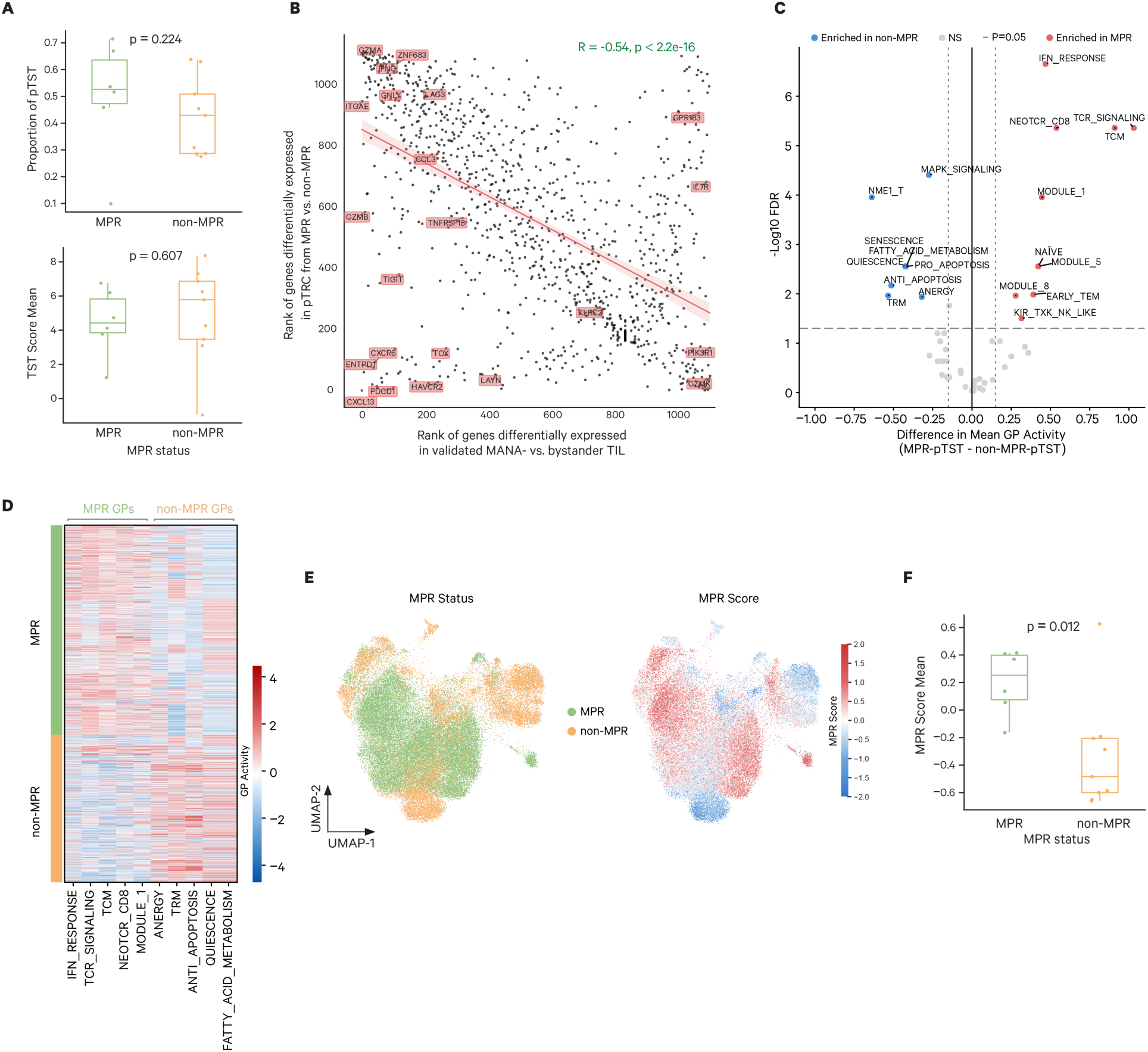
Functional pTSTs determines clinical response to immune checkpoint blockade (ICB). (**A**) Comparison of the mean proportion (top) and mean TST score (bottom) of intratumoral pTSTs between patients with major pathological response MPR (n=6) and non-MPR (n=9) in a NSCLC cohort. Statistical significance was determined by two-sided Mann-Whitney U tests, p*-*values are indicated. (**B**) Pearson correlation of ranks of differentially expressed genes (DEGs) between MPR-derived vs non-MPR pTSTs and validated MANA-specific vs bystander TILs. The correlation coefficient and p-value were labelled. (**C**) Volcano plot visualizes differential Gene Program activity between MPR and non-MPR groups, using a horizontal dashed line for the significance threshold (FDR<0.05) and vertical dotted lines to demarcate effect size boundaries (∣ΔZ∣>0.15) for enriched programs. (**D**) Heatmap illustrating the distinct activity profiles of the top five DGPs of pTST cells across MPR and non-MPR cell groups. (**E**) UMAP visualization of intratumoral pTSTs (n=56,885 cells) colored by the clinical response status (left) and the calculated MPR score (right). (**F**) Comparison of the mean MPR score in intratumoral pTSTs across MPR (n=6) and non-MPR (n=9) patients, statistical significance was determined by two-sided Mann-Whitney U tests, p-values are indicated.

We next analyzed the GP profiles of these intratumoral pTSTs to characterize their functional states across different response groups. Differential gene-program (DGP) analysis revealed marked differences in gene-program activity between MPR and non-MPR patients. pTSTs derived from MPR patients were preferentially enriched for gene programs associated with stemness (e.g., NAÏVE, MODULE_1 and MODULE_5) and early activation (e.g., TCR_SIGNALING, TCM and IFN_RESPONSE). By contrast, pTSTs from non-MPR patients exhibited elevated activity of gene programs typically enriched in TSTs, including those related to terminal exhaustion and dysfunction (e.g., NME1_T, ANERGY, PRO/ANTI_APOPTOSIS and SENESCENCE) (Fig. 5C).

Consistently, the differences in the activity of the top five DGPs between MPR-derived pTSTs and non-MPR-derived pTST enabled effective discrimination of pTSTs from the two patient groups (Fig. 5D). This critical biological distinction motivated the definition of the “MPR score”, a metric designed to quantify the functional potential of intratumoral pTSTs for evaluating MPR outcome. The MPR score was calculated as the summed activity of top five DGPs identified in pTSTs from MPR patients (IFN_RESPONSE, TCR_SIGNALING, TCM, NEOTCR_CD8, and MODULE_1). As expected, pTSTs derived from tumors of MPR patients exhibited higher MPR score (Fig. 5E). Moreover, the mean MPR score of intratumoral pTSTs at the individual patient level significantly discriminated patient groups with distinct responses to ICB therapy (p = 0.012) (Fig. 5F).

To further elucidate the biological interpretation of the MPR score, we examined its relationship with established stemness-related and Tpex-associated gene-expression signatures (fig. S4). Specifically, we compared the MPR score with a stemness enrichment score derived from canonical stemness markers (TCF7, IL7R and SELL)^17,37^, as well as three previously reported Tpex gene-expression scores defined in independent studies^38–40^. In contrast to the MPR score, none of these alternative scores consistently discriminated pTSTs derived from MPR versus non-MPR patients, nor did they distinguish patient groups at the individual-level (fig.S4A and B). Among the Tpex-related scores, the MPR score showed a significant positive correlation with the Tpex signature defined by Liu et al., but not with other Tpex scores (fig.S4C). By capturing Tpex-related features distinct from both canonical stemness and established transcriptional programs, the MPR score clarifies a unique functional state. Furthermore, these discrepancies emphasize the lack of consensus in defining the Tpex populations most relevant to successful immunotherapy outcomes.

### The MPR score is a robust and generalizable predictor of ICB response

To evaluate the robustness and generalizability of the MPR score in ICB response prediction, we applied the complete TSTScope pipeline, including pTST inference and MPR scoring, to an independent ICB-treated NSCLC cohort (n = 234) reported by Liu et al.^41^. Patients with fewer than 100 intratumoral CD8^+^ T cells were excluded to ensure reliable pTST inference. After quality control, 219,188 intratumoral CD8^+^ T cells from 221 patients with paired scRNA-seq and scTCR-seq data were retained for analysis (Methods). Using TSTScope, we identified 85,400 intratumoral CD8^+^ T cells as pTSTs in this cohort (fig. S5A). In the original study, Liu et al. partitioned CD8^+^ T cells into ten subsets and defined Tex-relevant cells as those sharing TCR clonotypes with terminally exhausted cells (CD8T_terminal_Tex_LAYN). Cells clonally linked to the terminal Tex subset but residing outside this cluster were annotated as Tpex. The pTSTs inferred by TSTScope showed extensive overlap with these Tex-relevant populations, predominantly mapping to CD8T_Tex_CXCL13 and CD8_Trm_ZNF683 subsets. Notably, pTSTs also included a substantial fraction of non-Tex relevant cells, mainly originating from CD8T_Trm_ZNF683, CD8T_Tm_IL7R, and CD8T_Tem_GZMK+GZMH+ cell subsets, with smaller contributions from CD8T_NK-like_FGFBP2 and CD8T_prf_MKI67 (Fig. 6A and B).

**Fig. 6.**
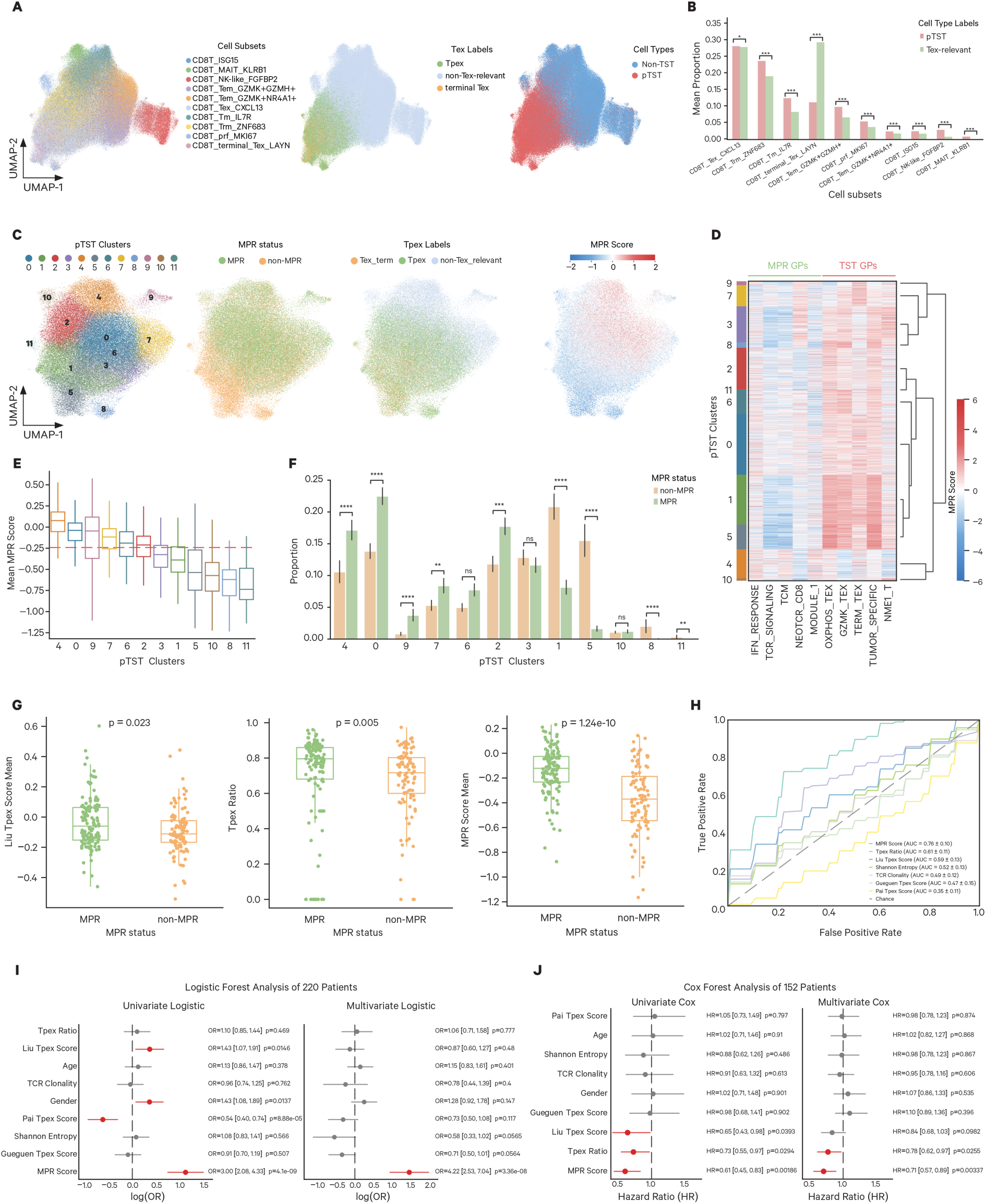
Robustness and generalizability of the MPR score in an independent validation cohort. (**A**) UMAP projections of CD8^+^ T cells (n=219,188) from Liu’s NSCLC cohort, colored by original cell subset annotations (left), study-defined Tpex/Tex labels (middle), and TSTScope-inferred pTST status (right). (**B**) Mean proportions of pTSTs (n=221 patients) and original study-defined Tex-relevant populations (n=215 patients) across identified cell subsets. Statistical significance was determined by two-sided Mann-Whitney U tests; p*-*values are indicated as *** *P* < 0.001 and * *P* < 0.05. (**C**) UMAP visualizations of sub-clustered pTSTs (n=85,400) identified by TSTScope, colored by cell clusters, clinical response status, original Tpex labels, and MPR score. (**D**) Heatmap plot of relative GP activities across pTST clusters, categorized by MPR-associated and TST-associated signatures. (**E**) Boxplot of MPR scores across cells in individual pTST clusters, ordered by mean score magnitude. The red dashed line denotes overall median score across pTST populations. (**F**) Comparison of per-patient pTST cluster proportions between MPR (n=123) and non-MPR (n=97) groups. Statistical significance was determined by two-sided Mann-Whitney U tests: \*\*\*\**P* < 0.0001, \*\*\**P* < 0.001, \*\**P* < 0.01, \**P* < 0.05; ns, non-significant. (**G**) Comparison of the patient’s Tpex-related metric (left), Tpex ratio (middle), and MPR score (right) between MPR (n=123) and non-MPR (n=97) groups. Two-sided Mann–Whitney U tests were used, p*-*values were indicated. (**H**) ROC curves evaluating the predictive performance of the MPR score versus Tpex- and TCR-based biomarkers for ICB response, AUC scores were labelled. (**I-J**) Forest plots depicting odds ratios (OR) from logistic regression (**I**) and hazard ratios from Cox proportional-hazards analysis (**J**) for the MPR score, alongside established ICB response biomarkers and clinical covariates. Results from univariate (left) and multivariate (right) analyses are shown with 95% confidence intervals and p-values.

To further resolve functional heterogeneity within pTSTs, we partitioned these cells into 12 subsets using the joint latent embedding learned by TSTScope. pTSTs from patients with distinct responses to ICB therapy displayed markedly different cluster preferences. Subsets enriched for non-Tex-relevant cells (C4, C7, and C9) were predominantly derived from MPR patients, whereas clusters characterized by terminal Tex features (C1, C5, C8, and C11) were largely contributed by non-MPR patients. Consistent with these compositional differences, pTSTs from MPR patients exhibited significantly higher MPR scores (Fig. 6C and fig. S5B). Moreover, MPR scores exhibited a continuous, graded distribution across pTST subsets rather than discrete stratification. These clusters exhibited hierarchical differences in the activity of MPR-associated and TST-associated gene programs: from C4, C0, and C2 to C1 and C5, decreasing MPR-program activity was accompanied by a progressive activation of TST-related programs (Fig. 6D). Clusters with higher MPR scores were enriched in MPR patients, whereas clusters with lower scores were predominantly derived from non-MPR patients (Fig. 6E and F).

We next benchmarked the predictive performance of the MPR score against established biomarkers, including TCR clonality, Shannon entropy^42,43^, three previously reported Tpex signatures, and the Tpex ratio defined by Liu et al.. Among these, only Liu’s Tpex signature score (p = 0.023) and the Tpex ratio (p = 0.005) were significantly elevated in MPR patients, whereas the MPR score demonstrated a far stronger association (p = 1.24 × 10⁻¹⁰) (Fig. 6G and fig. S5C). Receiver operating characteristic analysis further confirmed the superior predictive capacity of the MPR score (AUC = 0.76), outperforming the Tpex ratio (AUC = 0.61) and Liu’s Tpex signature score (AUC = 0.59) (Fig. 6H).

Consistent with these observations, logistic-regression analysis revealed a robust association between the MPR score and clinical response in both univariate (OR = 3.00, p = 4.1 × 10⁻⁹) and multivariate models (OR = 4.22, p = 3.26 × 10⁻⁸) (Fig. 6I). In a subset of patients with available follow-up data (n = 152), Cox proportional-hazards analysis demonstrated that higher MPR scores were significantly associated with improved recurrence-free survival (RFS) in both univariate (HR = 0.61, p = 0.00186) and multivariate settings (HR = 0.71, p = 0.00337) (Fig. 6J). Kaplan-Meier survival curves, stratified by the median MPR score, corroborated these findings; the high-score group exhibited significantly prolonged RFS compared to the low-score group. Notably, alternative metrics, including the Tpex ratio and previously established Tpex-based signatures, demonstrated weaker or non-significant prognostic value (Fig. S5D). Furthermore, comparative Cox forest plot analysis indicated that the MPR score achieved the highest prognostic concordance among all evaluated clinical and molecular features (Fig. S5E). Together, these data establish the MPR score as a robust and clinically informative predictor of response to immune checkpoint blockade (ICB) and long-term patient outcomes.

## DISCUSSION

The precise identification and functional characterization of tumor-specific T cells (TSTs) are paramount for deciphering the mechanisms of immune checkpoint blockade (ICB) and guiding next-generation immunotherapies. In this study, we present TSTScope, a multimodal deep learning framework that moves beyond simple data concatenation to create a biologically constrained, unified representation of T-cell identity. By effectively integrating transcriptional programs with TCR sequence features, our work yields critical methodological advances and a fundamental shift in understanding the determinants of ICB response.

First, we demonstrate that joint modeling conditioned by biological priors is essential for refining antigen-specific signals from noisy scTCR-seq data. A major challenge in current TCR analysis is that raw embeddings are often confounded by conserved germline V/J gene usage, which masks the critical antigen-specificity signals driven by the CDR3 region^44,45^. TSTScope addresses this by employing a gene program-constrained VAE that conditions TCR embeddings on the cellular transcriptional state. This strategy effectively filters technical noise and germline artifacts, producing representations that align more closely with antigen specificity than single-modality baselines or unconstrained models. Consequently, TSTScope enables the robust identification of neoantigen-specific T cells in an unsupervised manner, suggesting that multi-modal integration should become a standard for linking clonotype to function in the TME.

Second, applying TSTScope to ICB-treated NSCLC cohorts reveals a paradigm shift: the functional potential of TSTs, rather than their abundance, dictates therapeutic success. While previous strategies have focused on quantifying tumor-infiltrating lymphocytes or broadly reactive clones^10,46,47^, our analysis showed that the proportion of potential TSTs (pTSTs) in tumor tissue does not correlate with MPR. Instead, we identified a distinct functional dichotomy. pTSTs in non-responsive patients were characterized by terminal exhaustion and senescence programs. Conversely, pTSTs in responders exhibited a progenitor-like state enriched for early activation, TCR signaling, and stemness modules^35,37,48^. This specific functional phenotype captured by our defined “MPR score” suggests that the clinical benefit of ICB relies on a reservoir of TSTs that retain the plasticity to expand and differentiate, rather than the accumulation of terminally differentiated effectors.

The translation of these biological insights into the MPR score provides a robust clinical biomarker. In an independent validation cohort, the MPR score outperformed established biomarkers, including *CXCL13* expression, Shannon entropy, and notably, previously defined Tpex signatures. While the MPR score correlates with the Tpex concept, it proved superior to generic Tpex signatures in discriminating responders. This indicates that the “MPR state” represents a more specific functional niche, defined by the intersection of tumor-specificity and proliferative potential, that is uniquely critical for ICB efficacy. These findings underscore the utility of TSTScope not just for discovery, but for patient stratification, enabling a more granular assessment of the TME than “hot” versus “cold” classifications.

Our study has limitations that warrant further investigation. TSTScope is currently tailored to CD8^+^ T cells, incorporating CD4^+^ helper programs will be necessary to capture the full spectrum of anti-tumor immunity. Additionally, while the use of pre-defined gene programs ensures interpretability and robustness, it may limit the discovery of novel, uncharacterized biological pathways. Furthermore, high-quality paired scRNA-seq and scTCR-seq data remain a prerequisite, and while validated here in NSCLC, the universality of the MPR score across other cancer types requires confirmation.

In conclusion, TSTScope provides a high-definition map of T-cell heterogeneity, resolving the subtle interplay between antigen specificity and functional state. By shifting the focus from the quantity to the quality of tumor-specific T cells, we identify a decisive functional phenotype that predicts ICB response. These insights suggest that future therapeutic strategies should prioritize the preservation or expansion of this critical “MPR-positive” T-cell pool to improve clinical outcomes.

## MATERIALS and METHODS

### TSTScope Model Design

To jointly analyze single-cell gene expression and TCR sequence data while preserving biological interpretability, we developed TSTScope, a multi-modal Conditional Variational Autoencoder (CVAE) framework. TSTScope aligns scRNA-seq profiles and TCR embeddings into a shared, biologically meaningful latent space. The model comprises two parallel encoding branches, one for gene expression and another for TCR embeddings, projecting data into a shared latent space constrained by prior biological knowledge in the form of pre-defined T cell functional GPs (Fig. 1B).

### Gene Expression Module

For the transcriptomic modality, we implemented a CVAE architecture^22^ incorporating explainable constraints within the decoder to capture biologically interpretable latent representations. Let 𝑥_g_ ∈ 𝑅^𝐺^ denote the gene expression values for a cell, where G corresponds to the subset of genes annotated in at least one prior-defined GP. The encoder 𝐸_g_ compresses the high-dimensional input 𝑥_g_ into a low-dimensional latent space by estimating a mean vector and a variance vector σ_g_ . A probabilistic latent vector 𝑧_g_ ∈ 𝑅^𝐿^ is generated via reparameterization, where the *L* corresponds to the number of GPs derived from prior knowledge.

To enforce interpretability, the decoder 𝐷_g_ utilizes a structured constraint mechanism governed by a GP matrix 𝑀 ∈ {0,1}^𝐿×𝐺^ , a binary mask where the entry 𝑀_𝑖j_ = 1 indicates that gene *j* is annotated to Gene Program *i*, and 0 otherwise. The 𝑊_𝑑𝑒𝑐_ , are constrained via element-wise multiplication: 𝑊_𝑐𝑜𝑛𝑠𝑡𝑟𝑎𝑖𝑛𝑒𝑑_ = 𝑊_𝑑𝑒𝑐_ ⊙ 𝑀. This ensures that a latent GP variable contributes to a gene’s reconstruction only if biologically associated. The predicted mean expression 𝜇_𝑟𝑒𝑐_ is obtained by scaling the relative abundance with the total library size 𝑠 of each cell. The reconstruction is modeled using a Negative Binomial (NB) distribution as shown in equation (1).

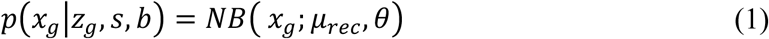

where 𝜇_𝑟𝑒𝑐_ = 𝑠 · softmax(𝑊_𝑐𝑜𝑛𝑠𝑡𝑟𝑎𝑖𝑛𝑒𝑑_^𝑇^𝑧_g_) represents the predicted mean expression adjusted by the library size 𝑠, and 𝜃 represents the gene-specific dispersion parameters conditioned on batch 𝑏. The reconstruction loss is defined as the negative log-likelihood of the gene expression counts, as represented in equation (2):

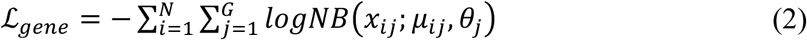

### TCR-Branch Module

We utilized the fine-tuned TCR-BERT model^21^ in inference mode to extract initial TCR embeddings denoted as 𝑥_𝑡𝑐𝑟_ ∈ 𝑅^𝐷^ . The TCR encoder, 𝐸_𝑡𝑐𝑟_is designed with an architecture analogous to the gene expression encoder. It accepts the initial embeddings 𝑥_𝑡𝑐𝑟_as input and projects them into a probabilistic latent space by estimating the Gaussian distribution parameters: a mean vector 𝜇_𝑡𝑐𝑟_ and a log-variance vector σ_𝑡𝑐𝑟_. Crucially, the dimensionality of this latent space is set to 𝐿, ensuring strictly consistent dimensionality with the gene expression latent space. A latent vector 𝑧_𝑡𝑐𝑟_ ∈ 𝑅^𝐿^ is subsequently obtained via reparameterization sampling. A symmetric Multi-Layer Perceptron (MLP) decoder, 𝐷_𝑡𝑐𝑟_ reconstructs the original TCR embeddings, with loss defined as Mean Squared Error (MSE) between the input embeddings and the reconstructed output, as shown in equation (3):

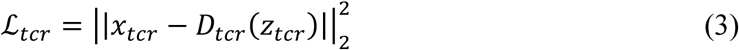

### Joint Latent Space Alignment and Fusion

To integrate the two modalities, we implemented a clonotype-guided alignment strategy that synchronizes the latent representations of the two modalities while preserving biological distinctiveness. We define a Center-Mean alignment loss 𝐿_𝑎𝑙𝑖g𝑛_that minimizes the distance between the centroids of cells sharing the same TCR clonotype across modalities. Let 𝐶 be the set of unique TCR clonotypes present in the current mini-batch. For each specific clonotype 𝑐 ∈ 𝐶, we compute the prototype (mean) latent vectors for both modalities as shown in equation (4):

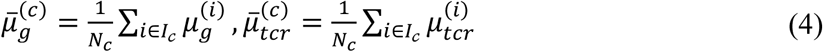

where 𝐼_𝒸_ denotes the set of indices for cells belonging to clonotype 𝑐, and 𝑁_𝑐_ is the number of such cells.

The alignment loss consists of three components, including inter-modality consistency by applying both Mean Squared Error (MSE) and Cosine Similarity loss between the paired Gene (𝜇̅^(𝑐)^) and TCR ( 𝜇̅^(𝑐)^) prototypes, and intra-modality distinctiveness by applying a contrastive loss specifically to the gene prototypes to encourage distinctiveness among different prototype representations. The total alignment loss is defined as in equations (5) and (6):

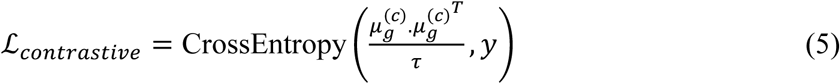

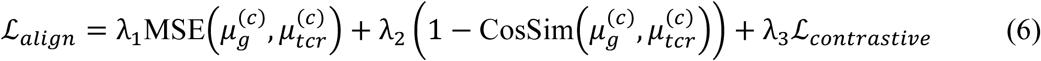

where 𝜏 is a temperature parameter and 𝑦 represents the identity labels for the clonotypes. Weights used in alignment loss were set to 𝜆_1_ = 1, 𝜆_2_ = 10, and 𝜆_3_ = 1. This objective pulls cells with identical TCRs towards the same functional state while ensuring that different clonotype prototypes remain distinguishable in the gene space.

Finally, a fused latent variable 𝑧_𝑓𝑢𝑠𝑖𝑜𝑛_ ∈ 𝑅^𝐿^ is computed via a linear combination of latent representation 𝑧_g_ and 𝑧_𝑡𝑐𝑟_, controlled by a fixed weighting hyperparameter λ = 0.5, followed by Layer Normalization.

### Joint Latent Space and Loss Function

The model is trained end-to-end by minimizing a composite loss function as in the equation (7):

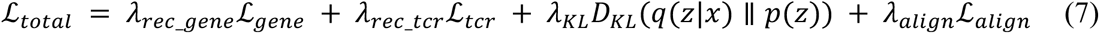

where ℒ_𝑁𝐵_ is the negative log-likelihood of the gene reconstruction, 𝐷_𝐾𝐿_is the Kullback-Leibler divergence regularizing the latent distributions towards a standard normal prior 𝑁(0, 𝐼), and ℒ_𝑎𝑙𝑖g𝑛_is the joint alignment loss. The weighting coefficients λ balance the reconstruction accuracy, latent space regularization, and multi-modal alignment. We used the Adam optimizer for model training. The hyperparameters used in this model listed in Table S2.

### Datasets and Preprocessing

**10x dataset**^49^: A publicly available dataset was obtained from the 10x Genomics website, comprising paired scRNA-seq and scTCR-seq profiles from four healthy donors, with TCR antigen specificities experimentally validated using peptide-MHC (pMHC) tetramers. From 33,804 CD8⁺ T cells from donor 2 annotated with TCR specificities spanning 29 antigen classes, the 6 antigen specificities with the largest clonal expansions were selected for antigen specificity classification, enabling evaluation of TSTScope’s capacity to capture antigen specific TCR features.

**Caushi et al. Cohort**^30^: This dataset includes paired scRNA-seq and scTCR-seq profiles from 16 anti-PD-1-treated patients with known MPR outcomes. Among these patients, 9 harbored experimentally validated neoantigen-specific T cells identified using MANAFEST. After quality filtering, the dataset comprised 188,804 CD8⁺ T cells across 88 samples, including 2,083 neoantigen-specific T cells (MANA) and 1,420 viral-specific T cells.

**Liu et al. cohort**^41^: This cohort comprises a comprehensive atlas of 1,254,749 tumor-infiltrating immune cells with paired scRNA-seq and scTCR-seq data from anti-PD-1 treated NSCLC patients. Based on the original cell-type annotations provided by Liu et al., 219,842 intratumoral CD8⁺ T cells were initially selected without additional quality-control filtering. Patients with fewer than 100 intratumoral CD8⁺ T cells were excluded. After filtering, 219,188 CD8⁺ T cells from 221 patients were retained for downstream analyses using the full TSTScope pipeline. Among these patients, 220 had available MPR status, and 152 had recurrence-free survival data.

**scRNA-seq Preprocessing**: scRNA-seq data generated using the 10x Genomics platform and preprocessed with Cell Ranger were analyzed using Scanpy (v1.8.1). Cells with fewer than 200 detected genes or more than 15% mitochondrial gene expression were excluded, and genes detected in fewer than 3 cells were removed. Quality-controlled expression counts were directly used as input for TSTScope, scAtlasVAE and ExpiMap. For PCA-based analyses, expression counts were further normalized to 10,000 transcripts per cell and log-transformed. Highly variable genes were identified by selecting the top 4,000 genes, with T cell receptor-related genes excluded based on IMGT annotation^50^ resulting in 3,894 genes retained for downstream analyses.

**scTCR-seq Preprocessing**: scTCR-seq data were obtained as the Cell Ranger processed contig annotation files. To ensure accurate matching between TCR sequences and antigen specificity in our benchmark dataset, cells expressing multiple TCRs were processed by retaining only the TCR chain with the highest UMI count as the representative sequence for each cell. scRNA-seq and scTCR-seq data were integrated using shared cellular barcodes, and only cells with available TCR β-chain CDR3 amino acid sequences were retained for downstream analyses and used as input for TSTScope.

### Method Comparisons and Implementation Details

We systematically benchmarked TSTScope against 6 representative single-modality feature embedding methods, encompassing state-of-the-art approaches for both TCR sequence and transcriptomic analysis.

**TCR sequence embedding models**: We evaluated 3 baselines for TCR representation learning:

**scCVC**^7^: We performed model inference using the scCVC foundation model. The TCR β amino acid sequences were extracted from the single-cell dataset and processed through an Embedding Wrapper utilizing a pre-trained Transformer checkpoint. For each sequence, we extracted features from the final hidden layer and applied a mean-pooling strategy to aggregate token-level information into a fixed-size embedding vector.

**TCR-BERT & Fine-tuned TCR-BERT**^21^: The TRB sequences were extracted from the single-cell metadata and processed through the TCR-BERT encoder with a pre-trained baseline version and a fine-tuned version optimized for specific downstream tasks. To capture deep contextual representations of the amino acid motifs. Using a layer-wise extraction strategy, features were retrieved from the final hidden layer to ensure the inclusion of the most refined semantic information. To condense the sequence-length outputs into cell-level representations, a mean-pooling method was employed, averaging the hidden states across all tokens to produce a fixed-dimensional latent vector for each cell.

**Transcriptomic feature extraction methods**: For gene expression analysis, we compared TSTScope against 3 established workflows:

**ExpiMap**^22^: We performed gene expression learning using the ExpiMap framework. Gene expression profiles were processed through a conditional variational architecture constrained by predefined GP gene sets (Table S1). To ensure a rigorous comparison, the model was configured using the same prior knowledge base integrated into TSTScope. The resulting latent embeddings represent biologically regularized features where each dimension corresponds to a specific functional program.

**scAtlasVAE**^32^: Utilizing a reference-mapping paradigm, transcriptomic profiles were projected into a high-dimensional latent space defined by a pre-trained reference model. We employed the human CD8^+^ T cell atlas as the structural scaffold for embedding inference. By aligning query cells to this established reference, we extracted fixed-size latent vectors that capture the relative positioning of individual cells within the broader immunological landscape.

**PCA with Harmony**^31^: Representing the scRNA-seq data integration baseline, we utilized identified HVGs to focus on the most informative transcriptomic features. Following Principal Component Analysis (PCA) to reduce dimensionality, we applied the Harmony algorithm to perform iterative batch correction. This strategy aligns cells across disparate donors and experimental conditions, yielding a corrected principal component space used as the final embedding representation.

### Gene Program Activity Quantification and Differential Analysis

To ensure the biological interpretability of the latent space, the sign of each gene program (GP) latent variable was aligned with its corresponding gene expression profile. For each GP, an aggregate expression score was calculated based on its constituent gene set. We then determined the Pearson correlation between this expression score and the corresponding latent dimension. To ensure that positive latent values consistently represent higher biological activity, latent values were multiplied by a direction factor derived from the sign of this correlation. These direction-corrected GP activity scores were then standardized across all cells using Z-score normalization to facilitate downstream visualization and comparative analyses.

Differential GP activity between cell populations was assessed using a non-parametric Bayesian framework. For each comparison, empirical distributions of GP activity were generated via bootstrapping (1,000 iterations per group). We estimated the posterior probability that activity in the target group (A) exceeded that in the reference group (B). The significance of this difference was quantified using the Bayes factor (BF), calculated as in equation (8):

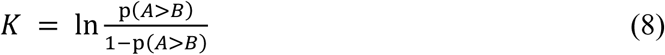

GPs were identified as significantly differentially active if the absolute Bayes factor exceeded a threshold of 1.1.

### Prediction and Evaluation of Tumor-Specific T cell Subpopulations

To quantify tumor specificity, we defined a TST score as the net difference between the aggregate expression of positive Gene Program (GP) markers (TUMOR_SPECIFIC, TERM_TEX, OXPHOS_TEX) and negative markers (EARLY_TEM, NAÏVE, TCR_SIGNALING).

The global distribution of TST scores across the dataset was modelled with a two-component Gaussian Mixture Model (GMM). To ensure a robust fit, we employed multiple random initializations and selected the final model based on the Bayesian Information Criterion (BIC). We assessed component separability by calculating the ratio of the distance between means to the sum of their standard deviations. The classification threshold was determined by the intersection of the two weighted Gaussian densities via numerical root-finding. In cases where an intersection was mathematically invalid or separability fell below a predefined heuristic, the global median score was used as the fallback threshold. Cells exceeding this threshold were labeled as potential TSTs (pTSTs), while the remainder were classified as non-TSTs. We validated the predictive accuracy of the TST score using MANA-specific T cells as the ground truth. Cells labeled as “undefined” were excluded from the validation cohort.

We further quantified and visualized clonotype composition and functional gene set enrichment profiles to characterize differences between pTST and non-TST groups. Clonal analysis was restricted to expanded clonotypes (size > 5), with log-transformed proportions calculated for group-wise comparisons. Functional module scores, including TRM, Checkpoint, Cytotoxicity, and TCR signaling signatures, were computed using the “score_genes” function in Scanpy. Statistical significance of enrichment differences was assessed using the Mann-Whitney U test. Differential gene expression (DGE) analysis was performed for both pTST versus non-TST and MANA versus Viral contrasts using the Wilcoxon rank-sum test with the “rank_genes_groups” function in Scanpy. To evaluate the consistency of transcriptional programs, concordance of gene rankings across comparisons was quantified using Pearson correlation coefficients with associated p-values and visualized via scatter plots.

### Benchmarking against Tumor-Reactive Signatures

To evaluate the predictive performance of TSTScope in identifying pTSTs, we compared the TST score against several established transcriptomic signatures validated for tumor-reactive CD8^+^ T cells. These benchmarks represent diverse approaches to defining tumor specificity, ranging from single-gene markers to large-scale transcriptomic programs.

**MANA score**^14^: Calculated following the methodology of Zeng et al., this score utilizes a weighted expression of CXCL13, ENTPD1, and IL7R to differentiate neoantigen-specific (MANA) CD8⁺ T cells from bystander populations.

**CXCL13 score**^10^: Evaluated as a high-resolution single-gene proxy for tumor reactivity by quantifying *CXCL13* expression at the single-cell level.

**NeoTCR8 score**^28^: A comprehensive 243-gene signature defined by Lowery et al., designed to capture the transcriptional landscape of neoantigen-reactive T cells derived from metastatic cancer cohorts, the signature scores were computed on raw expression matrices using gene set based scoring and used for downstream evaluation.

**Oliveira score**^27^: Computed using the gene set defined by Oliveira et al., representing transcripts significantly upregulated in tumor-specific CD8^+^ tumor-infiltrating lymphocytes (TILs) relative to virus-specific bystander T cells. The signature scores were computed on raw expression matrices using gene set based scoring and used for downstream evaluation.

### Formulation and Evaluation of the MPR Score

To quantify the therapeutic potential of the tumor-reactive T cell repertoire, we developed the Major Pathological Response (MPR) Score, a composite transcriptomic metric. The development and validation of this score proceeded through three main phases: feature identification, score formulation, and comparative benchmarking.

**Identification of MPR-associated Gene Programs:** To identify transcriptional programs associated with clinical response, single-cell GP activities were first aggregated to the patient level by calculating mean activity scores for each sample. We assessed differential GP activity between patients who achieved MPR and non-MPR using a two-sided Mann–Whitney U test. Gene programs demonstrating a statistically significant difference (P<0.05) were selected as candidate features for the final score.

**Formulation of the MPR Score:** The MPR score was designed to integrate multiple signals of effector potential into a single metric. For each patient-specific tumor-reactive T cell (pTST) population, the score was defined as the mean activity of the top-5 significantly positively enriched gene programs associated with treatment response, as identified in the preceding step. This approach allows for a robust representation of the functional state of the T cell repertoire rather than relying on individual gene expression.

**Benchmarking against Established Biomarkers:** To evaluate predictive performance, we benchmarked the MPR score against established transcriptomic signatures and repertoire-based metrics associated with tumor-reactive or progenitor exhausted (Tpex) CD8⁺ T cells.

**Liu et al. Tpex signature**^38^: Gene list derived from longitudinal single-cell profiling of NSCLC patients undergoing anti-PD-1 therapy, capturing a precursor exhausted T cell state characterized by stem-like and checkpoint-associated markers. The signature scores were computed on raw expression matrices using gene set based scoring and used for downstream evaluation.

**Gueguen et al. Tpex signature**^39^: Gene list describing a memory-like CD8⁺ T cell population with regenerative capacity identified in NSCLC. The signature scores were computed on raw expression matrices using gene set based scoring and used for downstream evaluation.

**Pai et al. Tpex signature**^40^: Gene list defined through multi-regional lineage tracing of tumor-specific T cells, marking a clonally expandable progenitor population persisting across lymphoid and tumor compartments. The signature scores were computed on raw expression matrices using gene set based scoring and used for downstream evaluation.

**TCR Repertoire Shannon entropy**^42^: Calculated from the frequency distribution of unique TCR clonotypes within each patient, as defined in Equation (9):

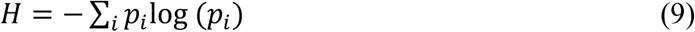

where 𝑝_𝑖_ denotes the relative abundance of clonotype 𝑖.

**TCR Repertoire Clonality**^42^: Defined as the complement of normalized Shannon entropy, as defined in Equation (10):

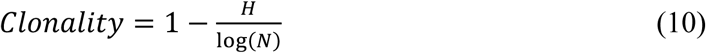

where *N* is the total number of unique clonotypes, yielding values ranging from 0 (fully polyclonal) to 1 (highly oligoclonal).

### Performance Evaluation of Patient ICB Response Clinical Outcomes

The predictive utility of the MPR score in the context of ICB therapy was assessed through a multi-level statistical framework designed to evaluate discriminative power, clinical independence, and longitudinal survival.

**Discriminative analysis**: Differences in MPR score distributions between responders and non-responders were evaluated using two-sided Mann-Whitney U tests. The score’s ability to differentiate clinical response was quantified using Receiver Operating Characteristic (ROC) curve analysis. Area Under the Curve (AUC) values were calculated from continuous score distributions using scikit-learn (v1.3.0), facilitating a direct comparison of the MPR score against established molecular benchmarks.

**Logistic Regression and Predictive Independence**: To assess whether the MPR score serves as an independent predictor of ICB response, we utilized univariate and multivariate logistic regression models via statsmodels (v0.14.0). Prior to modeling, all continuous predictors were Z-score standardized to ensure comparability of effect sizes. Multivariate models were adjusted for baseline clinical covariates, including age and sex, to account for potential confounding factors. Results are reported as Odds Ratios (ORs) with 95% Confidence Intervals (CIs).

**Survival analysis**: The association between MPR scores and clinical outcomes, specifically Recurrence-Free Survival (RFS), was evaluated using Cox proportional hazards regression implemented in lifelines (v0.28.0). Patients were stratified into high- and low-risk groups based on the median of MPR score, Tpex ratio, and Liu Tpex score. Survival trajectories were visualized using Kaplan–Meier curves, with differences between strata assessed via the log-rank test. To evaluate the global predictive accuracy of the survival models, we calculated Harrell’s Concordance Index (C-index), measuring the model’s ability to correctly rank order the time-to-event outcomes.

### Statistical Analysis

Statistical analyses were performed in Python 3.10.14, using functions in scikit-learn (v1.3.0), statsmodels (v0.14.0), and lifelines (v0.28.0) packages. Continuous variables were compared using the two-sided Mann-Whitney U test. For all frequentist tests, a p-value < 0.05 was considered statistically significant.

## Acknowledgments

We thank technical support from the Data Science Platform of Guangzhou National Laboratory and the Bio-medical Big Data Operating System (Bio-OS).

## Funding

National Key R&D Program 2023YFF1204701

Major Project of Guangzhou National Laboratory GZNL2025C01013, SRPG22007 National Natural Science Foundation of China 12371485, 82473482

Natural Science Foundation of Zhejiang Province Grants LY24H160007

Prevention and Control of Emerging and Major Infectious Diseases-National Science and Technology Major Project 2025ZD01901900

## Author contributions

Conceptualization: YXL, JWL

Methodology: JWL, SWC

Investigation: JWL, SWC, JYC

Visualization: JWL, SWC, JYC, FAW, CXY, JJC, KYW

Supervision: YXL, JWL, LLL

Writing-original draft: JWL, SWC, JYC

## Data and materials availability

The TCR antigen dataset is available from the 10x Genomics public repository (https://www.10xgenomics.com/datasets/cd-8-plus-t-cells-of-healthy-donor-2-1-standard-3-0-2). The dataset of ICB response cohorts from Caushi et al. and Liu et al. are available via the Gene Expression Omnibus (GEO) under accession numbers GSE173351 and GSE243013, respectively. The TSTScope software architecture was developed using the scArches framework^33^, and custom scripts used for data processing, model training, and analysis can be accessed on GitHub at https://github.com/YUZIXD/TSTScope. Processed data objects and integrated metadata required to replicate the findings of this study are available https://doi.org/10.5281/zenodo.18162425 under DOI 10.5281/zenodo.18162424.

