## Supplemental figures(fig.S1-fig.S5) and supplemental Tables(TableS1-TableS2)) for "TSTScope Unifies Single-Cell Multi-Omics to Identify Functional T Cell States Predictive of Immunotherapy Response"

Shiwei Cao *et al.*

;
.

**This PDF file includes:**

Figs. S1 to S5
Tables S1 to S2

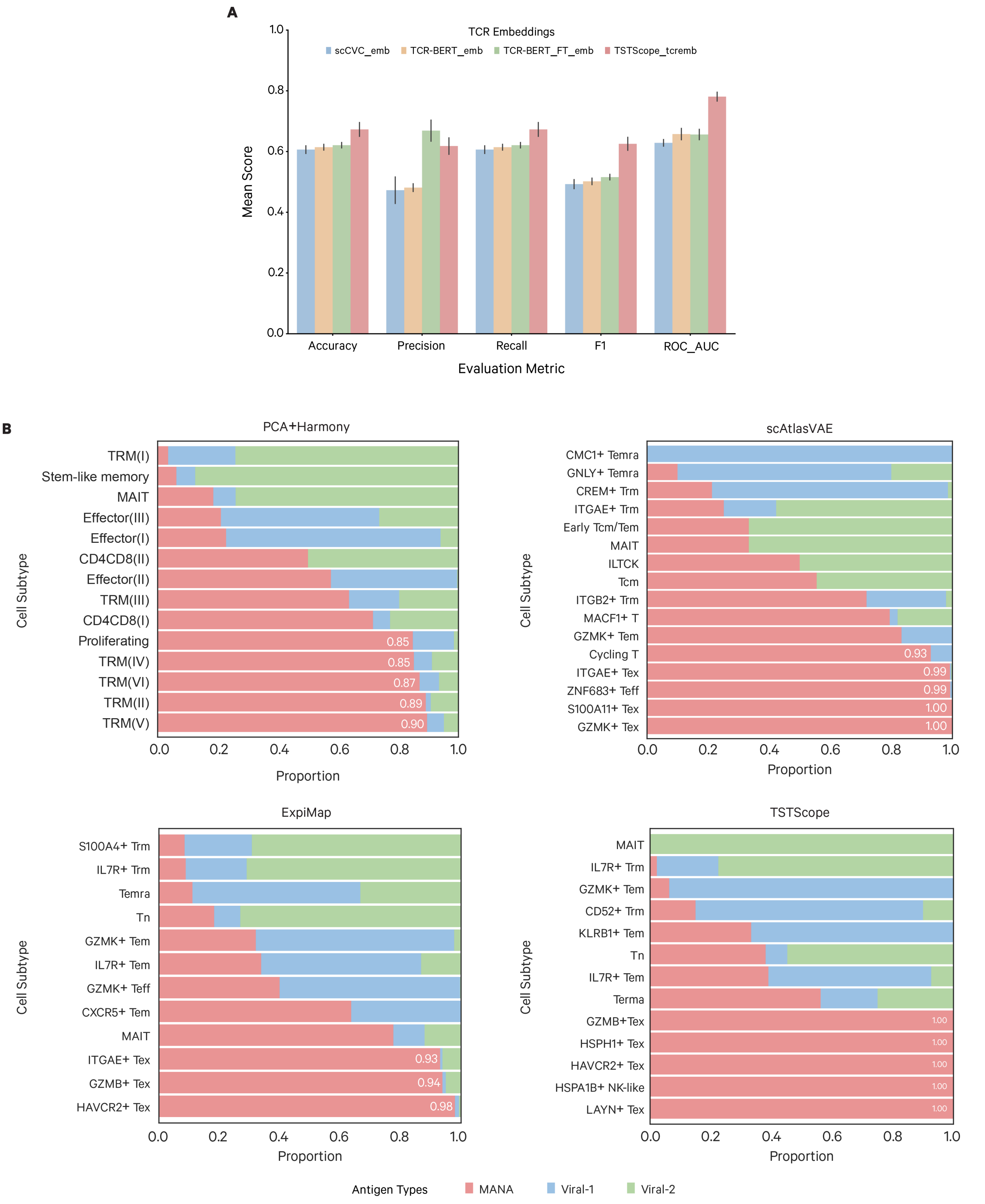


**Fig. S1 Benchmarking of TCR embeddings and antigen specificity distribution across T-cell subsets.** (**A**) Comparative performance of TSTScope and baseline TCR embeddings (scCVC, TCR-BERT, and TCR-BERT_FT) across five classification metrics. Data are presented as mean

± s.d. (**B**) Proportions of MANA (red), Viral-1 (blue), and Viral-2 (green) specific T cells within annotated subtypes across four approaches. T-cell subtypes with a MANA enrichment exceeding 85% are annotated with white numerals.

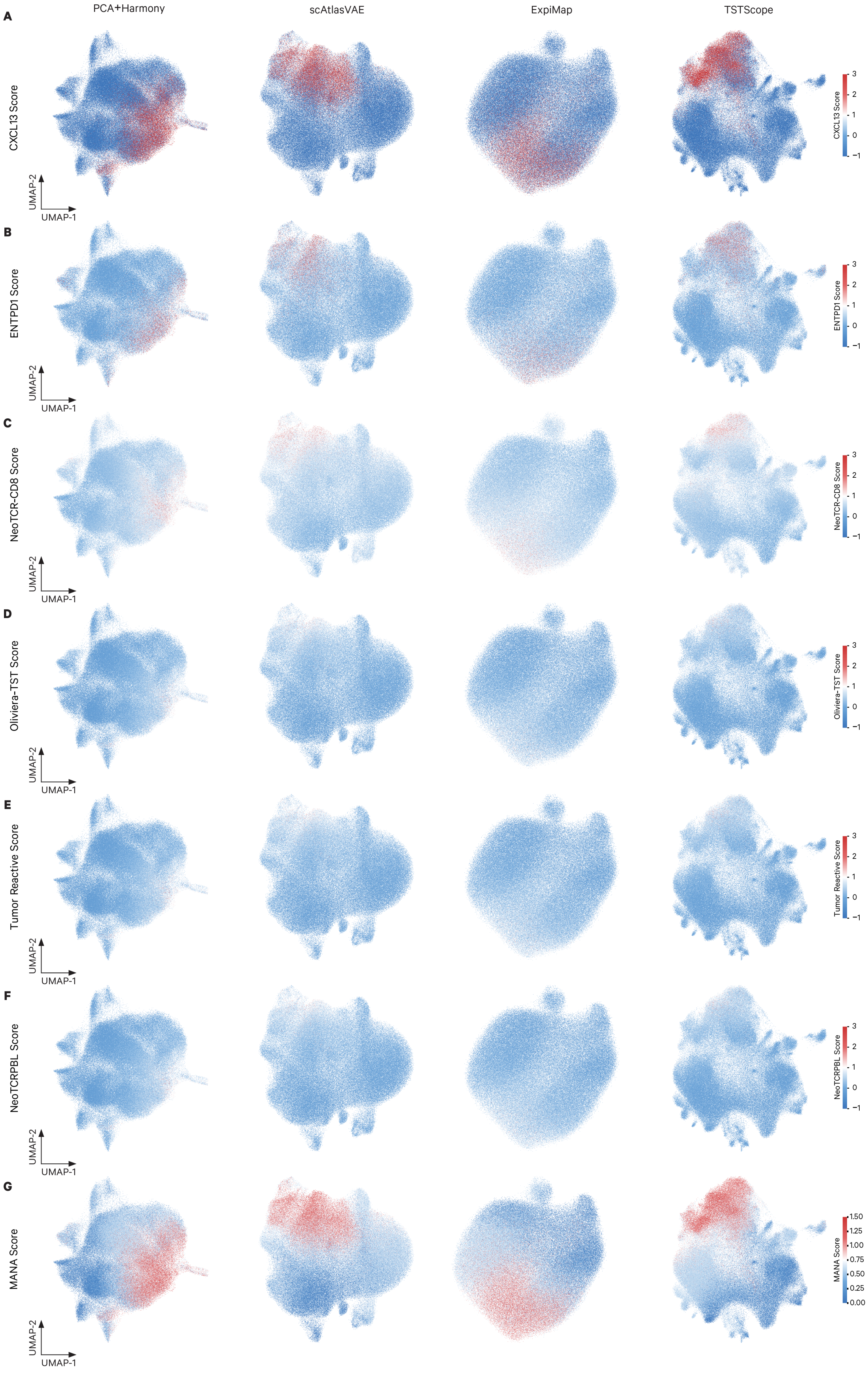

A
B
C
D
E
F
G
PCA+Harmony
scAtlasVAE
ExpiMap
TSTScope
CXCL13 Score
ENTPD1 Score
NeoTCR-CD8 Score
Oliviera-TST Score
Tumor Reactive Score
NeoTCRPBL Score
MANA Score
UMAP-1
UMAP-2
3
2
1
0
−1
1.50
1.25
1.00
0.75
0.50
0.25
0.00

**Fig. S2 Distribution of benchmark tumor-reactive signatures across latent integration methods.** (**A-G**) UMAP visualizations of 188,804 CD8$^+$ T cells from NSCLC patients (n=16) integrated using PCA+Harmony, scAtlasVAE, ExpiMap, and TSTScope, colored by normalized enrichment scores for established tumor-reactive markers and signatures: (**A**) CXCL13 expression; (**B**) ENTPD1 expression; (**C**) NeoTCR-CD8 signature; (**D**) Oliveira-TST signature; (**E**) Tumor Reactive signature; (**F**) NeoTCRPBL signature; and (**G**) machine learning-derived MANA score. Color scales represent relative signature activity or gene expression levels within the respective latent spaces.

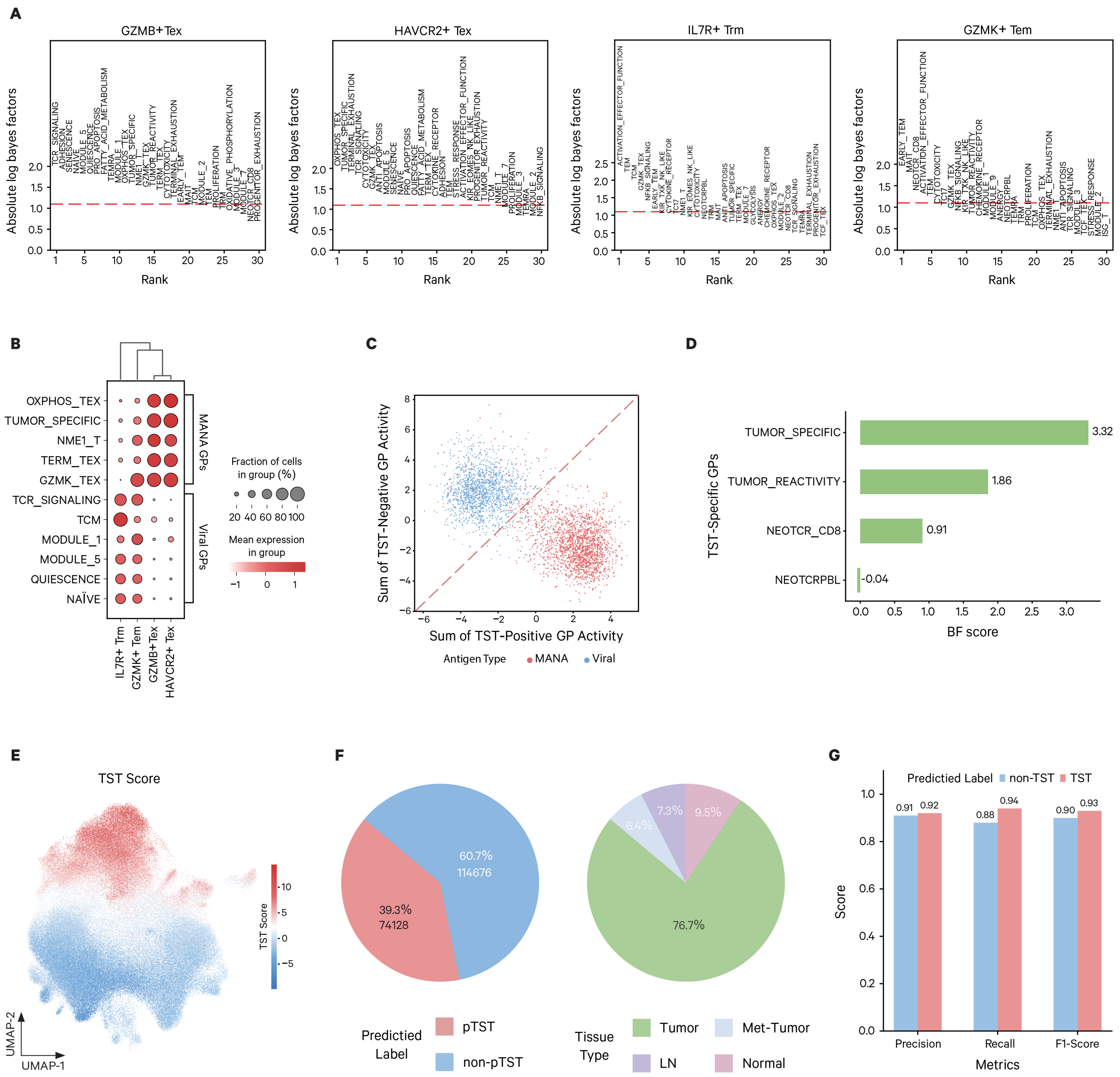


**Fig. S3 Characterization of latent gene programs and evaluation of pTST identification.** (**A**) Rank-ordering of gene programs by absolute log Bayes factors within representative TST-associated (GZMB+ Tex, HAVCR2+ Tex) and viral-associated (IL7R+ Trm, GZMK+ Tem) subsets. (**B**) Dot plot showing mean activity and fraction of cells for MANA- and Viral-associated gene programs across specific T-cell clusters. (**C**) Bivariate distribution of validated MANA-specific and Viral-specific cells based on cumulative activities of positive and negative TST gene programs. (**D**) Bayes factor (BF) scores for different tumor-specific gene programs. (**E**) UMAP plot of the NSCLC dataset (n=188,804 cells) colored by the calculated TST score. (**F**) Proportions of predicted pTST versus non-pTST labels (left) and the distribution of predicted pTSTs across different tissue origins (right). (**G**) Prediction performance metrics (precision, recall, and F1-score) for pTST and non-pTST identification based on ground-truth antigen-specificity labels.

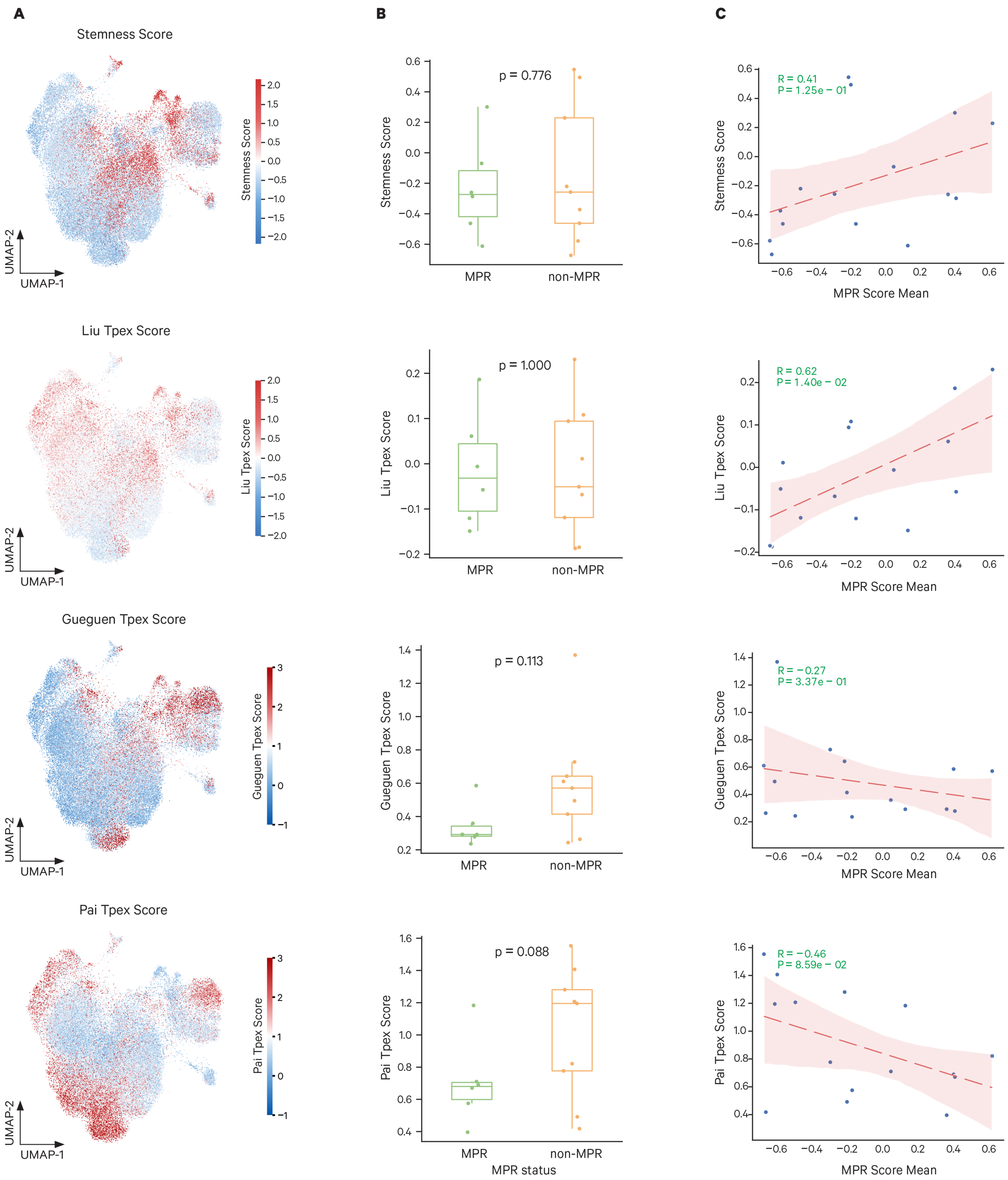


**Fig. S4 Relationship between the MPR score and established stemness or progenitor-exhausted T cell (Tpex) gene signatures.** (**A**) UMAP projections of intratumoral pTSTs (n=56,885 cells) colored by enrichment scores for a canonical stemness signature and three independently defined Tpex gene signatures (Liu *et al.*, Gueguen *et al.*, and Pai *et al.*). (**B**) Mean stemness and Tpex signature scores in intratumoral pTSTs from patients with MPR (n=6) and non-

MPR (n=9) in the NSCLC cohort. Statistical significance was determined by two-sided Mann-Whitney U tests, p-values were indicated. (**C**) Correlations between per-patient mean MPR scores and alternative stemness or Tpex signature scores across patients (n = 15). Pearson correlation coefficients (R) and corresponding p-values are indicated.

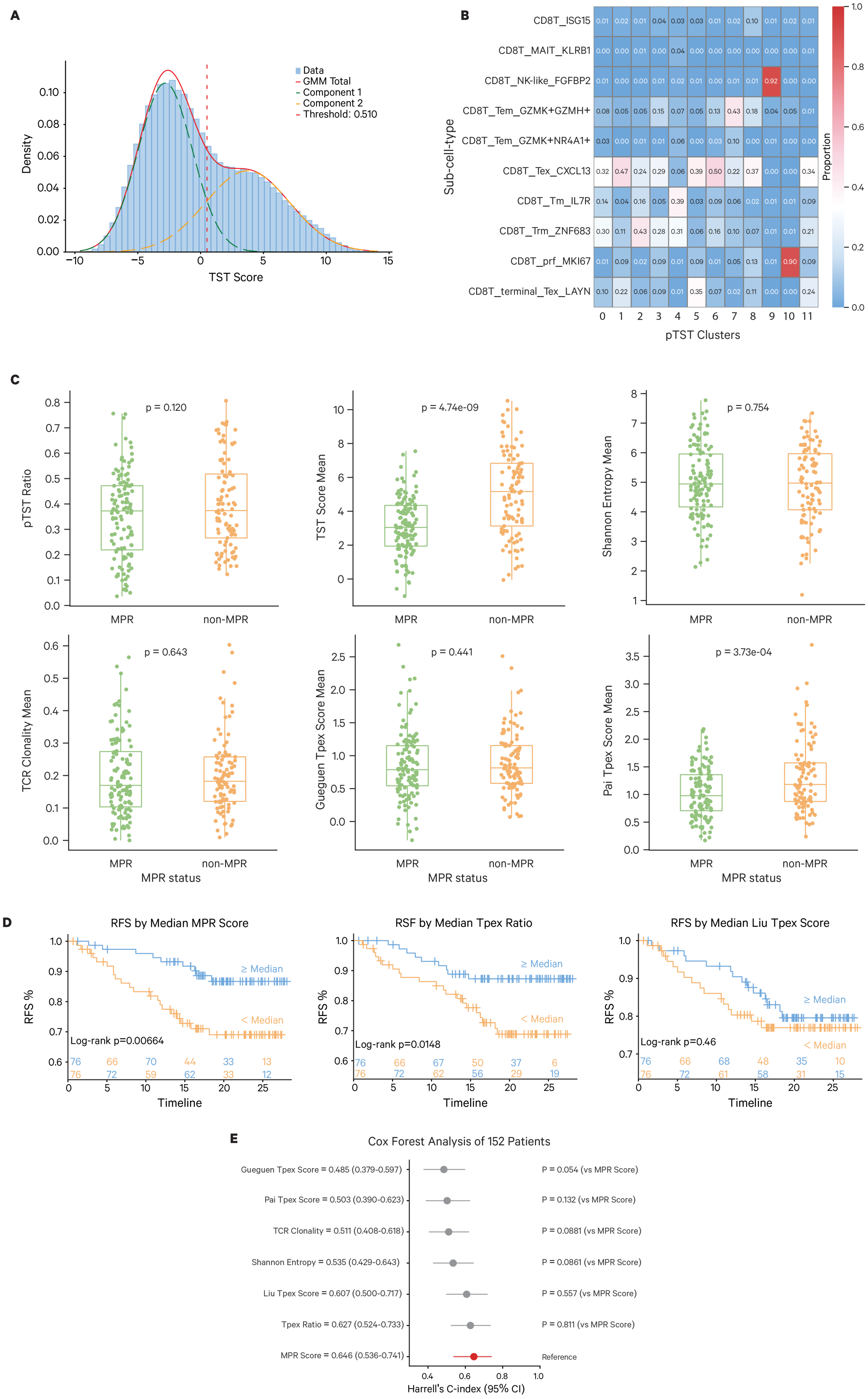
A
Data
GMM Total
Component 1
Component 2
Threshold: 0.510
Density
TST Score
B
CD8T_ISG15
CD8T_MAIT_KLRB1
CD8T_NK-like_FGFBP2
CD8T_Tem_GZMK+GZMH+
CD8T_Tem_GZMK+NR4A1+
CD8T_Tex_CXCL13
CD8T_Tm_IL7R
CD8T_Trm_ZNF683
CD8T_prf_MKI67
CD8T_terminal_Tex_LAYN
Sub-cell-type
pTST Clusters
Proportion
C
pTST Ratio
p = 0.120
TST Score Mean
p = 4.74e-09
Shannon Entropy Mean
p = 0.754
TCR Clonality Mean
p = 0.643
Gueguen Tpex Score Mean
p = 0.441
Pai Tpex Score Mean
p = 3.73e-04
MPR
non-MPR
MPR status
D
RFS by Median MPR Score
Log-rank p=0.00664
RSF by Median Tpex Ratio
Log-rank p=0.0148
RFS by Median Liu Tpex Score
Log-rank p=0.46
≥ Median
< Median
RFS %
Timeline
E
Cox Forest Analysis of 152 Patients
Gueguen Tpex Score = 0.485 (0.379-0.597)
Pai Tpex Score = 0.503 (0.390-0.623)
TCR Clonality = 0.511 (0.408-0.618)
Shannon Entropy = 0.535 (0.429-0.643)
Liu Tpex Score = 0.607 (0.500-0.717)
Tpex Ratio = 0.627 (0.524-0.733)
MPR Score = 0.646 (0.536-0.741)
P = 0.054 (vs MPR Score)
P = 0.132 (vs MPR Score)
P = 0.0881 (vs MPR Score)
P = 0.0861 (vs MPR Score)
P = 0.557 (vs MPR Score)
P = 0.811 (vs MPR Score)
Reference
Harrell's C-index (95% CI)

**Fig. S5 Robustness and prognostic utility of the MPR score in an independent validation cohort.** (**A**) Density distribution of the TST score modeled by the dual Gaussian mixture model (GMM) to define the classification threshold (0.510) for pTST identification in the Liu et al. cohort (n=221 patients). (**B**) Proportional distribution of TSTScope-defined pTST clusters across original study-defined $CD8^+$ T-cell subsets. (**C**) Comparison of per-patient mean values for the pTST ratio, TST score, TCR metrics (clonality and Shannon entropy), and Tpex signature scores (Gueguen *et al.* and Pai *et al.*) between MPR (n=123) and non-MPR (n=97) groups. Two-sided Mann–Whitney U tests were used, p-values are indicated. (**D**) Kaplan-Meier estimates of recurrence-free survival (RFS) stratified by the median scores of the MPR score, Tpex ratio, and Liu Tpex score (n=152 patients). (**E**) Comparative analysis of prognostic performance across various clinical predictors, ranked by Harrell's concordance index (C-index); p-values indicate statistical comparisons against the MPR score (red dot; reference) and 95% confidence intervals are shown.

### Supplementary Tables

**Table S1. Detailed description of the 50 curated T-cell Gene Programs (GPs).** This table provides a comprehensive summary of the 50 Gene Programs (GPs) used to constrain the TSTScope latent space, ensuring biological interpretability.

**Table S2. Hyperparameters and configuration for TSTScope model training.** This table details the specific hyperparameters and training configurations used to optimize the TSTScope multi-modal CVAE framework.

**Table S1. Detailed description of the 50 curated T-cell Gene Programs (GPs)**

| Signatures Types | Source of Signatures | Dim | Description of Signatures | Gene Sets |
|---|---|---|---|---|
| T cell regulation module | Zhou et al., Nature, 2023 | 0 | Module 1 | AR, ELF1, ERG, ETV6, FOS, HIPK2, LITAF, MAF, MAFF, NFE2, NFKB1, OTX1, PPARG, RFX1, RXRA, SMAD3, SOX5, SSBP3, ZNF362 |
| | | 1 | Module 2 | BTG2, ETS2, FOSL2, FOXL1, ID2, KLF4, LEF1, MAML2, NAB2, NPAS2, NR1H2, NRF1, PDLIM1, POU6F1, RREB1, TAL1, TFAM, TLE3, TRPS1, ZEB2 |
| | | 2 | Module 3 | BACH1, BATF, GATA3, HDGFL3, IKZF1, KLF8, MXD3, NCOA7, NFAT5, NFIL3, NR2F1, NR4A2, NR4A3, PRDM1, REL, RFX2, RUNX2, RUNX3, SREBF2, STAT4, TBX21, ZNF287, ZNF780A, ZHX2, ZSCAN12 |
| | | 3 | Module 4 | ATF6, BCL9, EGR1, EGR3, ELF2, EPAS1, FOSB, FOXN3, HIVEP1, ID3, JDP2, MAFK, MAX, MTA3, NR4A1, PBX3, RARA, RARG, RFX3, RFX5, RFX8, RUNX1, SOX4, SP1, SP3, SREBF1, ST18, SUB1, TSC22D1, ZBTB38, ZEB1, ZFP36L1, ZSCAN10 |
| | | 4 | Module 5 | AFF3, BACH2, BCL6, FOXP1, JUND, POU2F2, TCF12 |
| | | 5 | Module 6 | FHL2, FOXO1, GLIS1, IRF1, KLF5, NFIA, PCBP3, POU2AF1, PRKCB, RELA, RFX4, RORA, SMAD1, TCF4, TOX2, ZNF652 |
| | | 6 | Module 7 | ACTN1, EGR2, ETS1, FLI1, HMGB2, IRF7, MEF2A, MYB, MZF1, PLAGL1, RERE, SATB1, TCF7, TGIF1 |
| | | 7 | Module 8 | ARNT2, FOXB1, HIVEP3, IKZF2, KLF13, KLF3, MXI1, NFATC2, NR1H3, RBPJ, TOX, ZNF212 |
| | | 8 | Module 9 | ARID5A, BHLHE40, CREB3L2, CUX1, DBX1, E2F7, ELF4, ESR1, HBP1, HIVEP2, IRF2BP2, IRF8, JUN, MAFG, NCOR2, NFE2L2, SETBP1, SOX6, SP4, STAT3, XBP1, ZBTB20, ZNF219, ZNF629 |
| T cell functional signatures | Chu et al., Nat Med, 2023 & Yang et al., iMate 2024 | 9 | Naive | IL7R, CCR7, SELL, FOXO1, KLF2, KLF3, LEF1, TCF7, ACTN1, FOXP1 |
| | | 10 | Activation:Effector function | FAS, FASLG, CD44, CD69, CD38, NKG7, KLRB1, KLRD1, KLRF1, KLRG1, KLRK1, FCGR3A, CX3CR1, CD300A, FGFBP2, ID2, ID3, PRDM1, RUNX3, TBX21, ZEB2, BATF, IRF4, NR4A1, NR4A2, NR4A3, PBX3, ZNF683, HOPX, FOS, FOSB, JUN, JUNB, JUND, STAT1, STAT2, STAT5A, STAT6, STAT4, EOMES |
| | | 11 | Progenitor exhaustion | CTLA4, HNF1A, SLAMF6, CD8A, CD8B, TNF, ICOS, TNFSF14, SATB1, EOMES, ID3, TCF7, CXCR3, GZMK, SELL, IL7R, CCR7, CXCL10, CD44, BCL6, CD27, CD28 |
| | | 12 | Terminal exhaustion | CXCL13, ENTPD1, PRF1, GZMH, GZMB, GZMA, LYST, BATF, TIGIT, LAG3, HAVCR2, PDCD1, PRDM1, RBPJ, SLAMF7, CCL4, TNFSF10, TBX21, NFATC1, HIF1A, TNFRSF4, CXCR6, PTPN7, EWSR1, HLA-DPB1, HLA-DPA1, UBXN1, PSMB9, LY6E, CCNDBP1, IRF9, LCP2, CD38 |
| | | 13 | TCR Signaling | CALM1, CALM2, CALM3, CAST, CD247, CD3D, CD3E, CD3G, CSK, DOK1, DOK2, FYN, LCK, NFATC2, NFATC1, NFATC4, NFATC3, PLEK, PAG1, PTPN11, PTPN2, PTPN22, PTPN4, PTPN6, PTPN7, PTPRC, PTPRCAP, S100A10, S100A11, S100A13, S100A4, S100A6, ZAP70, DUSP1, DUSP2, DUSP4, DUSP5, DUSP10, LAT, PLCG1, PLCG2, PPP3CA, PPP3CC, FOS, FOSB, FOSL1, FOSL2, JUN, JUNB, JUND, NR4A1, NR4A2, NR4A3, BATF, IRF4, SH2D2A |
| | | 14 | Cytotoxicity | GZMA, GZMB, GZMH, GZMK, GZMH, GNLY, PRF1, IFNG, TNF, SERPINB1, SERPINB6, SERPINB9, CTSA, CTSB, CTSC, CTSD, CTSW, CST3, CST7, CSTB, LAMP1, LAMP3, CAPN2, TIA1 |
| | | 15 | Cytokine/Cytokine receptor | CSF1, IL10RA, IL16, IL17RA, IL18RAP, IL21R, IL2RB, IL2RG, IL32, IL9R, ADAM10, ADAM8, METRNL, CD70 |
| | | 16 | Chemokine/Chemokine receptor | CCR4, CCR5, CCR7, CXCR3, CXCR4, CXCR5, CXCR6, CCL3, CCL4, CCL4L1, CCL4L2, CCL5, CXCL13, CXCL8, XCL1, XCL2 |
| | | 17 | Anergy | NT5E, IZUMO1R, LAG3, NRP1, DGKA, CBLB, RNF128, ITCH, NFATC2, EGR2, EGR3, NR4A1, TOB1 |
| | | 18 | NFKB Signaling | NFKB1, NFKB2, NFKBIA, NFKBIB, NFKBIZ, CHUK, IKBKB, IKBKG, REL, RELA, RELB |
| | | 19 | Stress response | HIKESHI, P4HB, UBA52, MAPK1, RPA2, RPA3, TNRC6A, TNRC6B, TNRC6C, NUP210, RBX1, UBB, UBC, CREBBP, ETS1, LAMTOR1, EED, LAMTOR2, HSP90AA1, RBBP4, LAMTOR4, LAMTOR5, PSMB10, CDKN1A, ANAPC11, PSMA1, ATG4D, PSMA2, WIPI2, PSMA3, TFDP2, PSMA4, ATM, CYCS, PSMA5, WDR45, ANAPC16, PSMA6, EHMT1, PSMA7, EHMT2, TERF1, PSMC1, ATR, PSMC3, SOD1, PSMC4, PSMC5, PSMC6, PSME1, ATG3, NCF1, PSME2, ATG5, CHMP2A, CHMP2B, GSK3B, RPS19BP1, TPR, MDM4, ELOB, CDC26, ELOC, ST13, RPS6KA1, EEF1A1, RPS6KA3, CHMP4A, DNAJB1, CAMK2G, RPS27A, TXN2, PRDX1, PRDX2, DNAJB6, FKBP4, PRDX3, MAP2K3, MTMR14, ATG12, PRDX5, RHEB, PRDX6, GSTP1, ATG14, RELA, GPX1, VHL, HSPA1A, UBE2D1, TERF2IP, HSPA1B, UBE2D2, TXN, UBE2D3, HSPH1, CYBA, CDK4, HSF1, CDK6, MAPKAPK3, CITED2, MAP1LC3B, PSMD12, HIF1A, HMGA1, PSMD13, MINK1, PSMD14, HSP90AB1, PTGES3, CXCL8, HSBP1, NFKB1, UVRAG, CAT, PSMB1, CDKN2C, PSMB2, CDKN2D, FOS, PSMB3, CCS, JUN, GABARAPL1, GABARAPL2, PSMB6, KDM6B, PSMB8, PSMD2, TNIK, PSMB9, PSMD3, RANBP2, ATOX1, SEM1, PSMD6, GABARAP, PSMD7, PSMF1, CHMP3, STAT3, PSMD8, TINF2, PSMD9, CHMP7, CCAR2, HSPA4, YWHAE, BAG1, HSPA5, HSPA6, BAG3, DYNLL1, HSPA8, RB1CC1, BAG5, ANAPC5, DYNLL2, VCP, DNAJC2, CABIN1, CEBPB |
| | | 20 | MAPK Signaling | MAP2K1, MAP2K2, MAP2K3, MAP3K4, MAP3K5, MAP3K8, MAP4K1, MAP4K5, MAPK1, MAPK3, MAPK11, MAPK13, MAPK14, MAPK1IP1L, MAPKAPK3, MAPK8, MAPK9, MAP2K7, MAPK10, MAP3K7 |
| | | 21 | Adhesion | ITGA1, ITGA4, ITGAE, ITGAL, ITGAM, ITGB1, ITGB2, ITGB7, SELL, SELPLG, S1PR1, VCAM1, ICAM2, ICAM3 |
| | | 22 | IFN Response | IFIT1, IFIT2, IFIT3, IFIT5, STAT1, STAT2, MX1, IRF1, IRF4, IRF7, IRF8, IRF9, ISG15, ISG20, IFITM1, IFITM2, IFITM3, OAS1, OAS2, OAS3, JAK1, JAK2, SOCS1, SOCS3, TRIM14, TRIM21, TRIM22, APOL1, APOL2, APOL6, IFNGR1, GBP1, GBP2, GBP4, GBP5, GBP3, BST2, CMPK2, DDX58, DDX60, DDX60L, IFI30, IFI35, IFI44, IFI44L, IFI6, IFIH1, PARP10, PARP12, PARP14 |
| | | 23 | Oxidative phosphorylation | ATP1B3, ATP2A3, ATP2B1, ATP2B4, ATP5A1, ATP5B, ATP5C1, ATP5D, ATP5E, ATP5EP2, ATP5F1, ATP5F1A, ATP5F1B, ATP5F1C, ATP5F1D, ATP5F1E, ATP5G2, ATP5G3, ATP5H, ATP5I, ATP5IF1, ATP5J, ATP5J2, ATP5L, ATP5MC1, ATP5MC2, ATP5MC3, ATP5MD, ATP5ME, ATP5MF, ATP5MG, ATP5MPL, ATP5O, ATP5PB, ATP5PD, ATP5PF, ATP6AP1, ATP6AP2, ATP6V0B, ATP6V0C, ATP6V0D1, ATP6V0E1, ATP6V0E2, ATP6V1D, ATP6V1F, ATP6V1G1, ATP8B2, ATP8B4, COX16, COX17, COX4I1, COX5A, COX5B, COX6A1, COX6B1, COX6C, COX7A2, COX7B, COX7C, COX8A, CYC1, MT.ATP6, MT.ATP8, MT.CO1, MT.CO2, MT.CO3, MT.CYB, MT.ND1, MT.ND2, MT.ND3, MT.ND4, MT.ND4L, MT.ND5, MT.ND6, NDUFA1, NDUFA10, NDUFA11, NDUFA12, NDUFA13, NDUFA2, NDUFA3, NDUFA4, NDUFA5, NDUFA6, NDUFAB1, NDUFAF3, NDUFAF4, NDUFAF8, NDUFB10, NDUFB11, NDUFB2, NDUFB3, NDUFB4, NDUFB6, NDUFB7, NDUFB8, NDUFB9, NDUFC1, NDUFC2, NDUFS2, NDUFS3, NDUFS5, NDUFS6, NDUFS7, NDUFS8, NDUFV1, NDUFV2, SDHA, SDHB, TCIRG1, UQCRB, UQCRC1, UQCRC2, UQCRFS1, UQCRH, UQCRQ, SURF1, SURF4 |
| | | 24 | Glycolysis | SLC2A3, SLC2A8, PFKFB3, ALDOA, ENO1, GAPDH, GPI, PGAM1, PGK1, PKM, TPI1, GYG1, MDH1, MDH2, LDHA, LDHB, ADH5, IDH2 |
| | | 25 | Fatty acid metabolism | ACAA1, ACADM, ACADVL, ACSL5, ALDH16A1, ALDH9A1, ECH1, ECHS1, HADHA, HSD17B10, SLC27A2, DECR1, PEX16, SCP2 |
| | | 26 | Senescence | KLRG1, B3GAT1, TP53, MAPK14, MAPK8, MAPK1, CDKN2A, CD3G, CD3D, CD3E, CD247 |
| | | 27 | Pro-apoptosis | BCL2L11, BID, BIN1, BIN2, BAX, BAK1, CASP8, CASP3 |
| | | 28 | Anti-apoptosis | BCL11B, BCL2L1, BAG3, BAG4, BIRC3, GADD45B |
| | | 29 | Quiescence | BTG2, KLF2, FOXP1, RUNX1, MYC |
| | | 30 | Proliferation | MKI67, BIRC5, AURKB, UBE2C, DLGAP5, HJURP, CDC20, GTSE1, TPX2, CD4, CD8A, CD3G, CD3D, CD3E, CD247, CD8B |
| Tumor reactive signatures | Lowery et al., Science, 2022 | 31 | NeoTCR-CD8 (243 genes)-Lowery et al., Science 2022 | ATP10D, GZMB, ENTPD1, KIR2DL4, LAYN, HTRA1, CD70, CXCR6, HMOX1, ADGRG1, LRRN3, ACP5, CTSW, GALNT2, LINC01480, CARS, LAG3, TOX, PTPRCAP, ASB2, ITGB7, PTMS, CD8A, GPR68, NSMCE1, ABI3, SLC1A4, PLEKHF1, CD8B, LINC01871, CCL4, NKG7, CLIC3, NDFIP2, PLPP1, PCED1B, CXCL13, PDCD1, PRF1, HLA-DMA, GPR25, CD9, TIGIT, HLA-DRB5, SYTL3, SLF1, NEK1, CASP1, SMC4, TSEN54, PLSCR1, GNPTAB, HLA-DPB1, PLEKHA1, ARHGAP9, ALOX5AP, SH3BP1, NCF4, NELL2, GATA3, PPM1M, TNFRSF1A, AC022706.1, MCM5, HLA-DRB1, TNFSF10, TRIM21, HDLBP, ERN1, CALHM2, SASH3, ACTA2, MAST4, CAPG, MPST, IGFLR1, GZMA, CD27, ITGAE, SLA2, RHOC, COMMD8, MYO1G, SP140, PHPT1, CD2BP2, PLEKHO1, STAM, MRPL16, IL2RB, ID2, TESPA1, GOLGA8B, MIS18BP1, VAMP5, DAPK2, HLA-DPA1, TSG101, IL4R, CCND2, CTSC, TRAF3IP3, NLRC3, ORAI3, GNLY, MIR155HG, CARD16, CD82, ECH1, JAML, EEF1G, ETFB, DAXX, RBM4, HCST, RAB27A, YPEL2, CHST12, ARPC1B, PDIA4, PDIA6, AC243960.1, TBC1D10C, PTPN6, PYCARD, BST2, BTN3A2, MTG1, MLEC, DUSP4, GSDMD, SLAMF1, IFI6, PCID2, GIMAP1, ITGA1, CSNK2B, CDK2AP2, MYO1F, AC004687.1, PTTG1, APOBEC3C, TSPAN14, MOB3A, STXBP2, LCP2, PLA2G16, LINC00649, CST7, TADA3, SIT1, APOBEC3G, SUSD3, CD3G, ATP10D, GZMB, ENTPD2, KIR2DL5, LAYN, HTRA2, CD71, CXCR7, HMOX2, ADGRG2, LRRN4, ACP6, CTSW, GALNT3, LINC01481, CARS, LAG4, TOX, PTPRCAP, ASB3, ITGB8, PTMS, CD8A, GPR69, NSMCE2, ABI4, SLC1A5, PLEKHF2, CD8B, LINC01872, CCL5, NKG8, CLIC4, NDFIP3, PLPP2, PCED1B, CXCL14, PDCD2, PRF2, HLA-DMA, GPR26, CD10, TIGIT, HLA-DRB6, SYTL4, SLF2, NEK2, CASP2, SMC5, TSEN55, PLSCR2, GNPTAB, HLA-DPB2, PLEKHA2, ARHGAP10, ALOX5AP, SH3BP2, NCF5, NELL3, GATA4, PPM1M, TNFRSF1A, AC022706.2, MCM6, HLA-DRB2, TNFSF11, TRIM22, HDLBP, ERN2, CALHM3, SASH4, ACTA3, MAST5, CAPG, MPST, IGFLR2, GZMA, CD28, ITGAE, SLA3, RHOC, COMMD9, MYO1G, SP141, PHPT2, CD2BP3, PLEKHO2, STAM, MRPL17, IL2RB, ID3, TESPA2 |
| | Oliviera et al., Nature, 2021 | 32 | Tumor Specific-Oliviera et al., Nature, 2021 | KRT86, RDH10, TYMS, HMOX1, GNG4, CXCL13, AFAP1L2, ACP5, MYO1E, LAYN, TNS3, TNFSF4, AKAP5, HAVCR2, ENTPD1, SLC2A8, AC243829.4, ZBED2, MCM5, CAV1, GOLIM4, TRAV21, VCAM1, PON2, MTSS1, CD38, TRBV11-2, MS4A6A, TOX2, CSF1, GALNT2, FXYD2, PLPP1, LMCD1, MYL6B, LAG3, HLA-DRA, IGFLR1, CCDC50, CD27, KIAA1324, CDKN2A, CD70, ABHD6, CTLA4, PDCD1, GEM, NUSAP1, TOX, CXCR6, NMB, HOPX, CLIC3, INPP5F, SNAP47, TSHZ2, HLA-DMA, SIT1, HLA-DRB1, TUBB, PYCARD, ADGRG1, HLA-DQA1, PRF1, HLA-DPA1, PTMS, CKS1B, HIPK2, CHST12, LSP1, FAM3C, SLC1A4, NUDT1, DNPH1 |
| | Kunes et al., NBT, 2023 | 33 | Spectra tumor reactivity program | MKI67, TNFRSF18, ENTPD1, ITGAE, CXCL13, TNFRSF9, IFNG, GZMB, CCL3, GNLY, FASLG, CCR1, PRF1, GZMH, HAVCR2, SLA, BATF, SH2D2A, LCP2, GBP1, ITM2A, GZMA, RHOH, ZBTB32, DOK2, BTN3A2, LAX1, IL10RB, CIC, TESPA1, SEMA7A, LTA, GFI1, CD8B, TNFRSF4, GADD45G, FBXW7, STAT5, NFATC1, KRT86, CD38, NR4A3, GZMM, GALNT2, LCK, MTHFD2, RBPJ, LAG3, LAYN, LINC02195 |
| | Yossef et al., Can Cell, 2023 | 34 | NeoTCRPBL | PASK, MT2A, MT1E, S100A4, SELL, LIME1, DENND10, ALOX5AP, COTL1, ITM2A, MT1X, EIF3A, AQP3, SMC4, UBXN11, YWHAB, S100A11, S1PR4, MYO1G, LIMS1, CHN1, RBPJ, TMSB10, CRIP1, LEF1, ITGB1, TMEM123, CD52, GYPC, ANXA2, SELPLG, NPDC1, LGALS3, PAG1, ANXA5, UCP2, CORO1B, CD82, CD55, NEAT1, ITGB7, VIM, CDC25B, TLE5, KLF2, S100A10, S100A6, ACTG1, TPM4, ITGAE, MIR4435-2HG, CNN2, FYB1, HLA-DQB1, FXYD5, SESN3, HLA-DRB1, BIN1, ISG20, TIGIT, RHOG, MED15, RNASET2, BEX3, CAPZB, CORO1A, LIMD2, MT1F, PFN1, ACTB, SLC1A4, NAP1L4, LSP1, SUSD3, ICAM2, TNFRSF25, CYTOR, GPSM3, NR3C1, TTN, SPOCK2, GSTK1, CD5, PLSCR3, LTB, OCIAD2, LEPROTL1, PRDX1, NME2, NDUFA12, RAB37, SH3KBP1, NIBAN1, SPN, FKBP5, NDUFB9, SHMT2, ZYX, SMAP2, CSGALNACT1, TRADD, ATP5PD, HLA-DRB5, ELOVL5, ARHGDIB, CTLA4, R3HDM4, GPR171, RCSD1, MYL6B, TSPO, IFITM1, CAMK4, CD27, TKT, PPP2R5C, VSIR, CD7, PAICS, S1PR1, VOPP1, RASA3, MPRIP, MCUB, EMB, GPI, RASGRP2, MTERF4, DUSP16, SIT1, CARHSP1, ZMIZ1, GMFG, RAC2, PLEC, DGKA, SLC25A5, PFKL, OPTN, ETS1, MSC, ARID5B, HLA-DPA1, MFNG, P2RY8, SARAF, MAP3K1, TRAPPC5, AC119396.1, SLC16A3, CD3D |

| Signatures Types | Source of Signatures | Dim | Description of Signatures | Gene Sets |
|---|---|---|---|---|
| CD8+ T cell subsets signatures | Zheng et al., Science, 2021 | 35 | CD8_Tcm | CD62L, CCR7, CD27, CD28, GPR183, CD44, CXCR4, CXCR5, IL7R , CD45RO, DKK3, EOMES, GZMK, ZFP36L2, TSC22D3, ZFP36, BTG1, ANXA1, LMNA, CD55, TNFAIP3, FTH1, RGCC, PABPC1 |
| | | 36 | CD8_Trm | ZNF683, HOPX, ID2, ZFP36L2, RBPJ, CKLF, IL32, GZMB, XCL1, CCL5, GZMA, XCL2, CXCR6, CXCR3, CAPG, TMSB4X, S100A4, LGALS3, ACTB, SH3BGRL3, CD52, LGALS1, ITGA1, LDLRAD4, IL7R, PRDM1, TGFBR2, ITGA1, SIPR1, CCR7, SELL |
| | | 37 | CD8_early-Tem | EOMES, GZMK, CXCR4, CD74, CXCR5, CCR4, CD44, DUSP2, CMC1, CST7, DKK3, SH2D1A, ENC1, TRAT1, DTHD1, PIK3R1, CRTAM, CD69 |
| | | 38 | CD8_Tem | EOMES, SUB1, GZMK, GZMA, CCL5, GZMH, IL32, CCL4, CD74, CCR5, CXCR3, HLA-DRB1, HLA-DPA1, COTL1, HLA-DPB1, HLA-DQA1, HLA-DQB1, HLA-DRB5, ITM2C, CST7, APOBEC3G |
| | | 39 | CD8_Temra | ASCL2, KLF2, KLF3, ZEB2, TBX21, GZMH, GZMB, GZMM, GZMA, CX3CR1, CXCR2, CMKLR1, CXCR1, FGFBP2, FCGR3A, S1PR5, PRSS23, GNLY, NKG7, KLRD1, FGR, PLEK, C1orf21 |
| | | 40 | CD8_TCF+ Tex | TSHZ2, TCF7, NR3C1, TOX, BATF, CXCL13, EBI3, TNFSF8, CD40LG, CCR7, IL6R, IFNAR2, CCR4, LHFP, CD200, GNG4, TNFRSF4, IGFL2, CPM, NMB, SESN3, BTLA, IGFBP4 |
| | | 41 | CD8_GZMK+ Tex | TOX, CXCL13, TCF7, SELL , IL7R, CCR7, GZMA, GZMK, CD74, CD200, CXCR5, CD27, CD28, EOMES, BATF, TNFSF8, CD40LG, CCR4, IL6R, TNFRSF4, ENTPD1, ITGB1, CCL3, CCL5, CCL3L3, TNFSF4, CCL4, IFNG, CD74, CXCR6, CCR5, HAVCR2, VCAM1, PDCD1, DUSP4, CTLA4, TNFRSF9, HLA-DQA1, HLA-DRB1 |
| | | 42 | CD8_term-Tex | RBPJ, ETV1, TOX, ZBED2, TOX2, CXCL13, TNFSF4, FAM3C, GZMB, CSF1, CCL3, CD70, IFNG, NAMPT, FASLG, IL2RA, CXCR6, CD74, IL2RB, IL2RG, TNFRSF9, LAYN, ENTPD1, HAVCR2, CTLA4, KRT86, TNFRSF18, GEM, TIGIT, DUSP4 |
| | | 43 | CD8_OXPHOS- Tex | ETV1, PRDM1, CXCL13, CSF1, CCL3, FAM3C, LAYN, HAVCR2, CTLA4, ENTPD1, TNFRSF9, TNFRSF18, PHLDA1, ACP5, TIGIT, KIR2DL4 |
| | | 44 | CD8_NME1+ T | ENO1, HMGA1, GTF3A, MYB, MXD3, HMGB1, MIF, GPI, CKLF, CD74, CXCR3, ACTB, MND1, SPC24, PFN1, ACTG1, CENPA, KIFC1, SKA3, GAPDH, KIF20A |
| | | 45 | CD8_MAIT | SLC4A10, IL4I1, CCR6, KLRB1, NCR3, LTB, RORA, IL18RAP, ZBTB16, IL23R, CD40LG, RORC, ADAM12, IL7R, CCL20, ZFP36L2, DUSP1, IFNGR1 |
| | | 46 | CD8_Tc17 | RORC, ZBTB16, CEBPD, RORA, NR1D1, CCL20, CD40LG, LTB, TNFSF13B, IL26, IL17A, FLT3LG, TNF, IL23A, CCR6, IL23R, IL7R, IL17RE, IL18RAP, SLC4A10, KLRB1, TMIGD2, IL4I1, NCR3, LTK, CA2, ME1, AQP3, ADAM12 |
| | | 47 | CD8_KIR+EOMES+ NK-like | IKZF2, EOMES, NR4A2, LITAF, ZNF331, GZMK, XCL2, XCL1, GZMM, TNFSF9, IFNGR1, CXCR4, CD74, IL2RB, KIR2DL3, KIR3DL2, CD160, TYROBP, KLRF1, KLRD1, KLRG1, CMC1, GCSAM, DUSP2 |
| | | 48 | CD8_KIR+TXK NK-Like | IKZF2, HOPX, TXK, FOSL2, REL, XCL1, XCL2, AREG, FAM3C, IL32, IL2RB, IL12RB2, KIR2DL3, KIR3DL2, KIR2DL4, KIR2DL1, KIR3DL1, TYROBP, KLRC2, KLRC3, LAT2, B3GNT7 |
| | | 49 | CD8_ISG+ T | PLSCR1, STAT1, IRF7, STAT2, SP100, TNFSF10, CCR1, CD74, IFIT1, RSAD2, IFIT3, IFI44L, MX1, IFI6, OAS1, CMPK2, ISG15, OAS3 |

**Table S2. Hyperparameters and configuration for TSTScope model training**

| Hyperparameter | Value | Description |
|---|---|---|
| Dropout Rate | 0.05 | Dropout probability for regularization |
| Gene Reconstruction Weight ($\lambda_g$) | 1 | Weight for the Negative Binomial loss |
| TCR Reconstruction Weight ($\lambda_{tcr}$) | 0.001 | Weight for the TCR reconstruction MSE loss |
| Alignment Weight ($\lambda_{align}$) | 1 | Weight for the joint latent space alignment loss |
| KL Divergence Weight ($\lambda_{KL}$) | 0.5 (Annealed) | Max weight for KL divergence, annealed from 0 to 0.5 over first 50 epochs |
| Optimizer | Adam | Optimization algorithm |
| Learning Rate | 1.00E-03 | Initial learning rate |
| Epsilon ($\varepsilon$) | 0.01 | Epsilon parameter for Adam optimizer |
| Weight Decay | 0 | L2 penalty for regularization |
| Batch Size | 512 | Number of cells per training batch |
| Epochs | 100 | Maximum number of training epochs |
| Random Seed | 2024 | Seed for reproducibility |
